# Ependymal cell maturation is heterogeneous and ongoing in the mouse spinal cord and dynamically regulated in response to injury

**DOI:** 10.1101/2022.03.07.483249

**Authors:** Aida Rodrigo Albors, Gail A. Singer, Andrew P. May, Chris P. Ponting, Kate G. Storey

## Abstract

The spinal cord neural stem cell potential resides within the ependymal cells lining the central canal. These cells are, however, heterogeneous, and we know little about the biological diversity this represents. Here we use single-cell RNA-sequencing to profile adult mouse spinal cord ependymal cells. We uncover transcriptomes of known subtypes and a new mature ependymal cell state, that becomes more prominent with age. Comparison of ependymal cell transcriptomes from the brain and spinal cord revealed that ongoing cell maturation distinguishes spinal cord ependymal cells from their postmitotic brain counterparts. Using an *ex vivo* model of spinal cord injury, we show that ependymal cell maturation is reversible but also highly regulated. We revisit ependymal cell identities in adult human spinal cord and uncover evidence for their maturation and surprising ventralisation with age. This first in-depth characterisation of spinal cord ependymal cells paves the way to manipulation of distinct ependymal subtypes, provides insights into ependymal cell maturation dynamics and informs strategies for coaxing ependymal cell-driven spinal cord repair.

## Introduction

A final fate of embryonic radial glial cells is to give rise to adult neural stem cells and ependymal cells (Cebrian Silla et al., 2021; Ortiz-Álvarez et al., 2019). Ependymal cells are ciliated cells lining the brain ventricles and the central canal of the spinal cord. Their functions, although largely speculative, include roles in barrier formation, as well as transport of ions, small molecules, and water between the cerebrospinal fluid (CSF) and the parenchyma (Bruni, 1998; Meunier et al., 2013). In the brain and spinal cord, ependymal cells are a key component of the subventricular zone and central canal stem cell niches (Becker et al., 2018; Hugnot and Franzen, 2011; Riquelme et al., 2008; Sabelström et al., 2013). While it is well-established that in the mouse brain ependymal cells are postmitotic and the neural stem cell potential resides in subependymal type B1 cells (Capela and Temple, 2002; Chiasson et al., 1999; Doetsch et al., 1997, 1999; Shah et al., 2018; Spassky et al., 2005), ependymal cells in the adult spinal cord can be found proliferating occasionally and at least some retain neural stem cell potential. From axolotls and zebrafish to mice, spinal cord ependymal cells can re-enter the cell cycle to self-renew and give rise to specialised cell types in response to injury and in culture (Barnabé-Heider et al., 2010; Bauchet et al., 2013; Dromard et al., 2008; Johansson et al., 1999; Lacroix et al., 2014; Li et al., 2018; Llorens-Bobadilla et al., 2020; Meletis et al., 2008; Mothe et al., 2011; Pfenninger et al., 2010; Stenudd et al., 2022). Ependymal cells in the human spinal cord have not been observed proliferating (Alfaro-Cervello et al., 2014; Dromard et al., 2008; Paniagua-Torija et al., 2018), but also display neural stem cell features when cultured *in vitro* (Dromard et al., 2008; Hugnot, 2013; Mothe et al., 2011). Despite the potential of spinal cord ependymal cells, the ability to repair the injured spinal cord varies greatly between species and spinal cord injuries most often result in permanent disability in mice and humans (Becker and Becker, 2015; Becker et al., 2018; Tazaki et al., 2017). Interestingly, while axolotls and zebrafish grow continuously and retain their regenerative ability throughout life (Holder et al., 1991), the proliferative capacity of ependymal cells in rodents declines with age (Alfaro-Cervello et al., 2012; Gonzalez-Fernandez et al., 2017; Li et al., 2018; Sabourin et al., 2009), in parallel with their *in vitro* neural stem cell potential (Li et al., 2018). Not surprisingly, ependymal cells have attracted the attention of researchers interested in harnessing this endogenous cell population to promote spinal cord repair (Barnabé-Heider and Frisén, 2008). However, a closer look at the central canal in mice and humans suggests that ependymal cells are heterogeneous, and we know surprisingly little about what this heterogeneity reflects.

Morphologically, spinal cord ependymal cells in the mouse are usually classified in three subtypes: radial, cuboidal, and tanycytes; even though cells with intermediate shapes are common (Alfaro-Cervello et al., 2012; Bruni and Reddy, 1987; Meletis et al., 2008). Based on the expression of a handful of molecular markers, spinal cord ependymal cells are also molecularly heterogeneous (Alfaro-Cervello et al., 2012; Ghazale et al., 2019; Meletis et al., 2008; Petit et al., 2011; Sabourin et al., 2009). Importantly, whether this heterogeneity reflects different functions, different maturation states or both remains unknown. Human spinal cord ependymal cells appear similarly heterogeneous and undergo dramatic changes with age. While ependymal cells in infants and teenagers resemble those of the mouse and organise around the central canal, in adult humans the central canal tends to collapse into an ependymal cell mass (Alfaro-Cervello et al., 2014; Ghazale et al., 2019; Kasantikul et al., 1979; Milhorat et al., 1994; Torrillas de la Cal et al., 2021; Yasui et al., 1999). A limitation of many human studies is low sample sizes and so, how human and mouse ependymal cells align and the molecular changes that human ependymal cells undergo with age are poorly understood.

In this study, we use single-cell transcriptomics to dissect ependymal cell heterogeneity in the adult mouse spinal cord, uncovering the transcriptomes of known and novel ependymal cell subtypes and states. We discover a striking tardiness in the maturation of lateral ependymal cells that distinguishes these cells from their postmitotic brain counterparts. In an *ex vivo* spinal cord assay we demonstrate that ependymal cell maturation is dynamically lost but also rapidly re-gained in response to injury. Informed by our mouse data, we revisit ependymal cell identities in the adult human spinal cord and uncover evidence for their maturation and surprising ventralisation with age.

## Results

### A cell map of the spinal cord central canal region

To comprehensively characterise spinal cord ependymal cells, we took two complementary single-cell RNA-sequencing (scRNA-seq) approaches: droplet-based 3′ scRNA-seq (10x Genomics), to profile a large number of cells in and around the central canal; and plate-based full length scRNA-seq (Smart-seq2), to profile ependymal cells in depth. Using the 10x approach, we profiled cells in the central canal region from 3-month-old wild-type mice (n = 4) (Figure 1A). After aggregating the four scRNA-seq libraries and filtering out low-quality cells and doublets we retained 9,061 cells, with a median of 7,631 transcripts (unique molecular identifiers or UMIs) and 2,672 features detected per cell (Figure S1A). For the plate-based scRNA-seq dataset, we isolated GFP-positive ependymal cells from age-matched FOXJ1-EGFP reporter mice (Ostrowski et al., 2003) (n = 4) by fluorescence activated cell sorting (FACS) and used Smart-seq2 technology (Picelli et al., 2013) to generate a full-length scRNA-seq dataset of 904 high-quality ependymal cell transcriptomes, with a median of 997,488 sequencing reads and 3,844 features detected per cell (Figure S2A).

**Figure 1.**
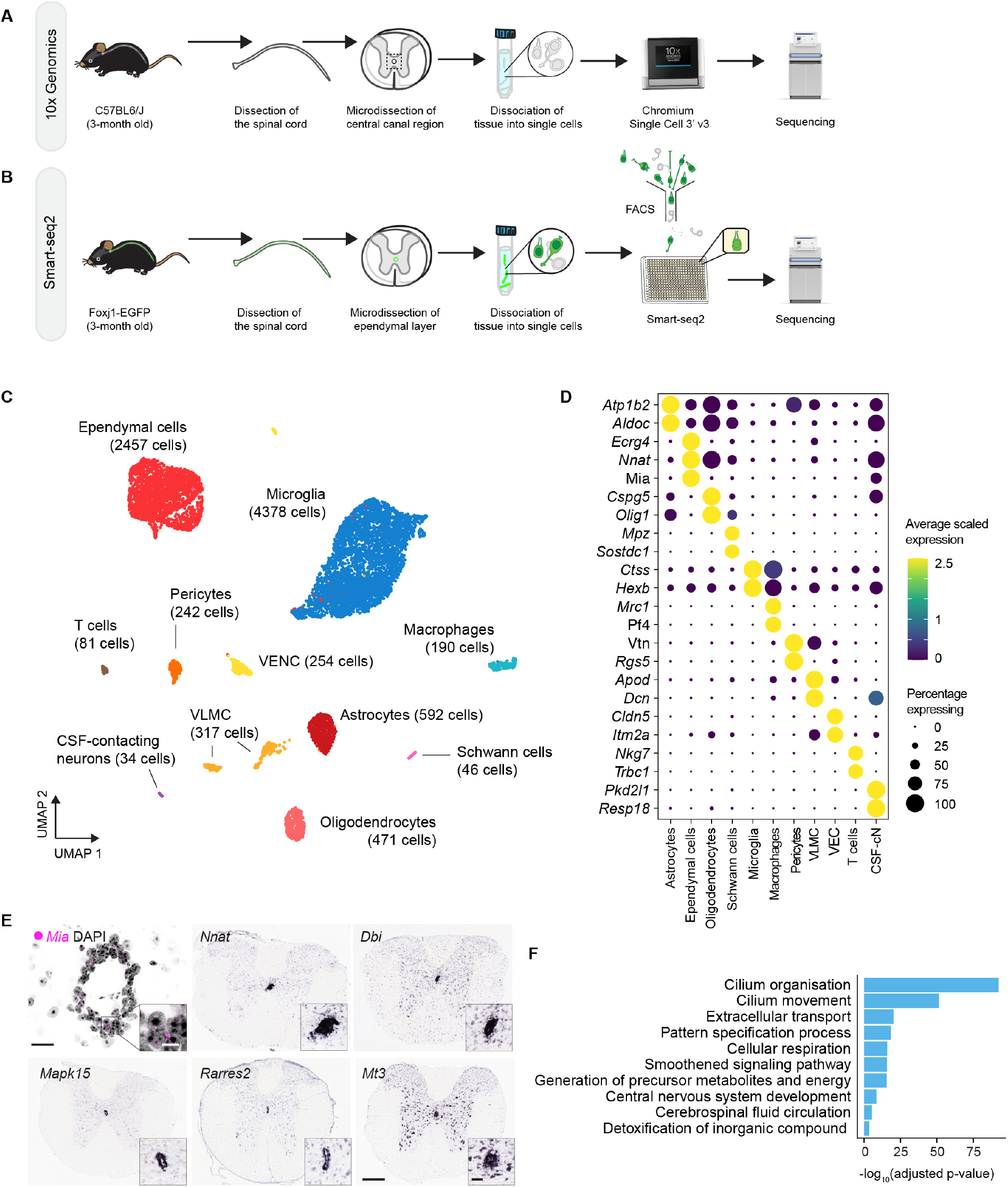
A cell map of the spinal cord central canal region. (A) Workflow for the generation of the 10x scRNA-seq dataset. (B) Workflow for the generation of the Smart-seq2 scRNA-seq dataset. (C) UMAP embedding of cells from the central canal region that passed QC (D) Dot plot of selected marker genes for each cell type in (C) and the percentage of cells within each cluster expressing each gene. (E) Close-up of the central canal region showing RNAscope for *Mia* (magenta dots) and *in situ* hybridization images from the Allen Brain Atlas showing the expression of ependymal-enriched genes *Nnat*, *Dbi*, *Mapk15*, *Rarres2*, and *Mt3*. Insets in images from the Allen Brain Atlas show a close-up view of the central canal. Scale bars: 30 μm and 20 μm (insets) (RNAscope images); and 200 μm and 50 μm in (Allen Brain Atlas images). (F) Selected GO terms associated with genes whose expression is enriched in ependymal cells. Bars show biological processes significantly overrepresented ranked by adjusted p-value (false discovery rate or FDR).

We first focused on the broader 10x dataset to explore the unique molecular features of ependymal cells compared to that of other cell types in the central canal region. After normalising the data, we identified highly variable features and used these for clustering using Seurat’s graph-based clustering approach (Butler et al., 2018; Stuart et al., 2019). The cells partitioned into 27 clusters (Figure S1B) which we manually annotated into 11 major cell types based on the expression of known marker genes (Figure 1C and 1D, Table S1). Microglia was the most abundant cell type represented in the dataset, followed by ependymal cells, oligodendrocyte-lineage cells, astrocytes, vascular leptomeningeal cells, vascular endothelial cells, pericytes, macrophages, T cells, and CSF-contacting neurons. All clusters contained cells derived from all samples, which suggests that the clustering was driven by biological heterogeneity rather than by sample-to-sample or technical variability (Figure S1C).

Differential expression analysis identified *Ecrg4* and *Mia* as two of the most specific and highly expressed genes in ependymal cells (Figure 1D and Table S1). Both genes encode secreted factors that inhibit proliferation of neural stem cells (Gonzalez et al., 2011; Kujuro et al., 2010; Nakatani et al., 2019) and neuroectodermal tumour cells (Hau et al., 2002), respectively. We confirmed the ependymal-specific expression of *Mia* using RNAscope technology (Figure 1E). This supports the notion that ependymal cells are secretory cells that play a key role in regulating neural stem cell niches and CNS homeostasis. We also confirmed that the expression of some ependymal-enriched genes is consistent with *in situ* hybridisation data from the Allen Brain Atlas (http://mousespinal.brain-map.org/) (Figure 1E). To infer further biological functions of ependymal cells we performed Gene Ontology (GO) enrichment analysis on the full list of ependymal-enriched genes (Figure 1F). Unsurprisingly, genes associated with cilium-related terms were overrepresented, consistent with the role of ependymal cilia contributing to CSF flow. The GO terms extracellular transport and detoxification of inorganic compounds were also significantly overrepresented, highlighting ependymal regulation of CSF composition. Other overrepresented biological processes were cellular respiration and CNS development (Table S2).

### Ependymal cells in the mouse spinal cord are transcriptionally heterogeneous

Lineage tracing studies and the expression of progenitor-specific transcription factors indicate that spinal cord ependymal cells derive from three embryonic neural progenitor domains: the roof plate (Shinozuka et al., 2019; Xing et al., 2018), the floor plate (Canizares et al., 2019; Ghazale et al., 2019; Khazanov et al., 2017), and the p2 and pMN domain (Fu et al., 2003; Ghazale et al., 2019; Masahira et al., 2006; Yu et al., 2013). However, surprisingly little is known about the molecular makeup and functions of these distinct ependymal cell subtypes.

To explore comprehensively ependymal cell heterogeneity, we subset the ependymal cell cluster from the 10x dataset and integrated it with the Smart-seq2 dataset using canonical correlation analysis (CCA) (Stuart et al., 2019) (Methods). The integrated dataset consisted of 2,792 ependymal cells with a median of 11,821 UMIs/reads and 3,479 features detected per cell (Figure 2B and Figure S2A and B). Clustering of the cells revealed six ependymal cell subtypes (Figure 2A). Nearly all samples from both technologies contributed cells to each of the six clusters (Figure S2C). Consistent with their different developmental origins, we found that all ependymal cell subtypes expressed one of the three sets of domain-specific markers (Figure 2C). Dorsal ependymal cells were clearly identified by expression of the roof plate marker *Zic1* (Nagai et al., 1997) while ventral ependymal cells expressed *Arx*, a well-known floor plate marker (Miura et al., 1997). The rest of ependymal cell clusters co-expressed the transcription factors *Pax6* and *Nkx6-1*, similarly to p2 and pMN progenitors, and due to their location within the central canal (Fu et al., 2003; Ghazale et al., 2019) we refer to these cells as lateral ependymal cells.

**Figure 2.**
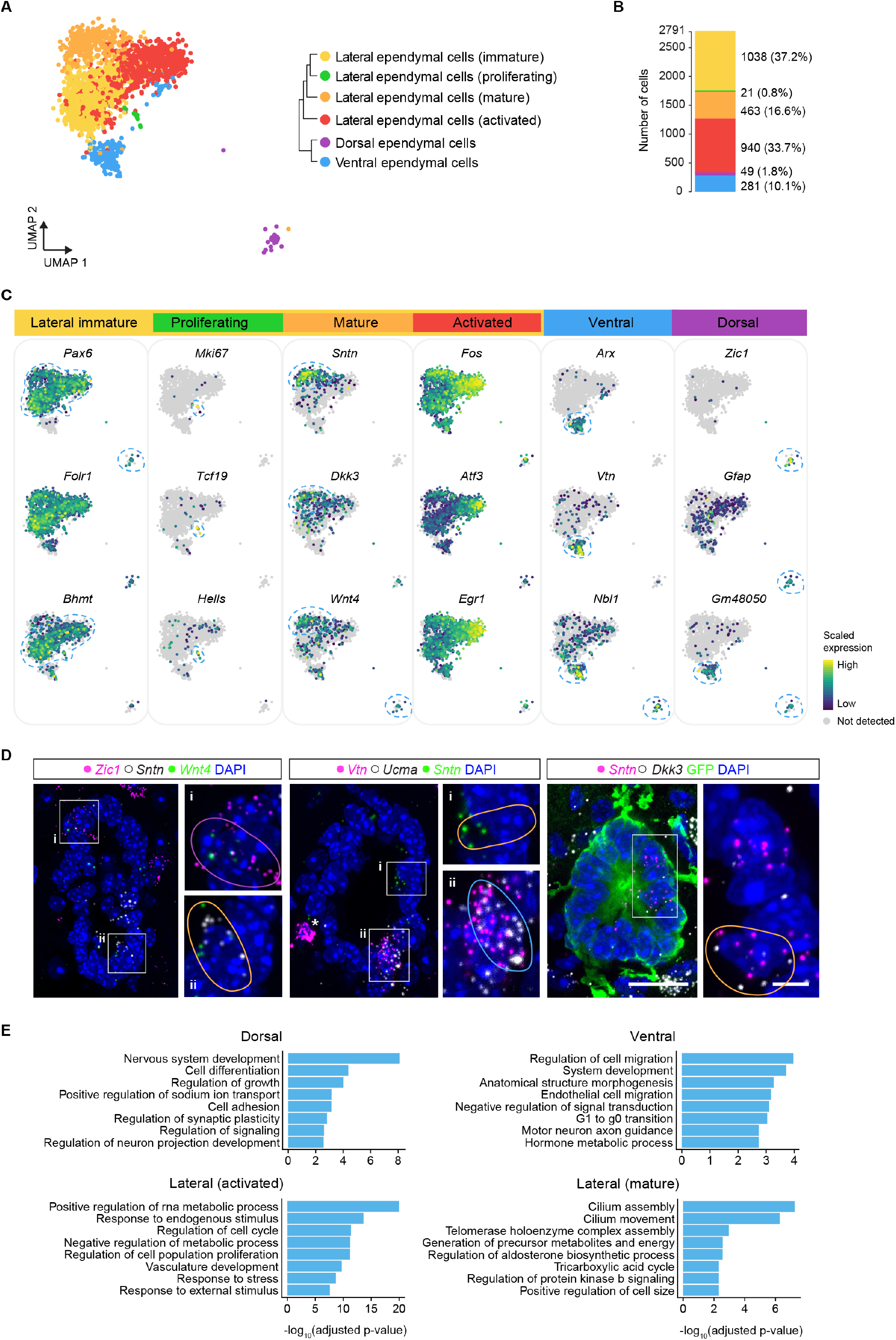
A cell map of spinal cord ependymal cells. (A) UMAP embedding of spinal cord ependymal cells captured using 10x and plate-based Smart-seq2 scRNA-seq technology. Each cell is coloured by their assigned ependymal subtype or state. Dendrogram shows the relationship between ependymal cell cell subtypes and states. (B) Bar plot showing the percentage of each ependymal cell subtype and state in the integrated dataset. (C) UMAP plots showing the expression distribution of selected marker genes. (D) RNAscope images confirming subset-specific expression of key marker genes. Cells representative of distinct subtypes are delineated and colour-coded by the colour assigned to them in (A). Scale bars: 20 μm and 5 μm (higher magnification images). (E) GO analysis of genes enriched in each ependymal cell subtype and state. Bars show biological processes significantly overrepresented ranked by adjusted p-value (FDR).

To identify genes further defining each ependymal subtype and to explore heterogeneity within lateral ependymal cells, we carried out differential expression analysis and used the list of genes enriched in each subtype to perform GO analysis and infer their functions (Table S3). Dorsal ependymal cells were characterised by expression of Zic transcriptions factors (*Zic1*, *Zic2*, *Zic4*, and *Zic5*) and expressed the highest levels of the intermediate filament genes *Gfap* and nestin (*Nes*) (Figure 2C and 2D). The abundant intermediate filaments are thought to confer robustness to the cytoskeleton of dorsal cells (Shinozuka and Takada, 2021; Sturrock, 1981). Dorsal ependymal cells also expressed genes encoding signalling molecules such as *Bmp6* and *Wnt4* (Figure 2C and 2D), supporting the notion that dorsal ependymal cells continue to act as a signalling centre in the adult spinal cord. Biological processes associated with dorsal cells include nervous system development, cell differentiation, regulation of growth, regulation of signalling, and regulation of synaptic plasticity (Figure 2E). Indeed, cellular compartment terms associated with dorsal cells suggest close association of these cells with neurons (e.g. integral component of postsynaptic density) (see Table S4 for full list of GO terms associated with each ependymal subtype). Together, these data suggest that dorsal ependymal cells retain some functions of embryonic radial glial cells such as providing structural support and contributing to the signalling microenvironment in the central canal niche. Dorsal ependymal cells also appear to share some functions with astrocytes such as monitoring and altering synaptic function, and likely modulate synaptic transmission in the dorsal spinal cord.

At the opposite pole of the central canal, ventral ependymal cells express floor plate genes such as the axon guidance cue *Ntn1* (Kennedy et al., 1994), the secreted bone morphogenetic protein (BMP) inhibitors *Nbl1* and *Ucma* (Surmann-Schmitt et al., 2008; Willems et al., 2018) (Figure 2C and 2D), and *Sulf1*, an extracellular heparan sulfate endosulfatase that modulates cell signalling (Dhoot et al., 2001; Ramsbottom et al., 2014). A high number of gene products expressed by ventral cells locate to the extracellular matrix and extracellular space, denoting the secretory nature of these cells. GO analysis revealed that among the molecular functions associated with ventral cells are glycosaminoglycan binding, growth factor binding, and metal chelating activity; and associated biological processes include regulation of cell migration, system development, and negative regulation of signal transduction (Figure 2E and Table S4). Taken together, this evidence suggests that ventral ependymal cells are a specialised ependymal cell subtype and a main contributor of cell extrinsic factors to the niche, including extracellular matrix components and signalling modulators. Interestingly, we noticed a few genes uniquely expressed by dorsal and ventral ependymal cells. These include the axon guidance cue, *Slit2*, and the unannotated gene *Gm48050* (Figure 2C). Dorsal and ventral ependymal cells could thus share some functions such as providing axon guidance cues postnatally and structural support to the central canal niche.

Lateral ependymal cells co-expressed *Pax6* and *Nkx6-1* and partitioned into 4 cell states (Figure 2A). An intriguing subset was enriched in cilia-related genes and uniquely expressed *Sntn*, a gene encoding for the apical ciliary protein Sentan (Kubo et al., 2008) (Figure 2C and 2D, Table S4). *Sntn* is a known marker of mature ciliated cells in the airways and oviducts (He et al., 2020; Konishi et al., 2016; Kubo et al., 2008; Mali et al., 2018), which suggests that it marks mature ependymal cells. Other molecular features of *Sntn* cells such as higher expression of the cyclin-dependent kinase inhibitor p21 gene (*Cdkn1a*) and histone genes (*Hist1h1c*, *Hist1h4h* and *Hist1h4i*), suggest ongoing change in the cell cycle state and chromatin organisation of these cells. Intriguingly, *Sntn* cells expressed the secreted factor *Dkk3,* a negative regulator of Wnt signalling (Zhu et al., 2014), and *Wnt4* (Figure 2C and 2D), encoding a ligand involved in non-canonical Wnt signalling that induces quiescence in the muscle stem cell niche (Eliazer et al., 2019). Moreover, *Sntn* cells also expressed *Phlda3,* a p53-regulated repressor of Akt and potential tumour suppressor (Kawase et al., 2009; Ohki et al., 2014). Together, this uncovers a unique signalling landscape within *Sntn* lateral cells. Besides cilium-related terms, other biological processes overrepresented in *Sntn* cells include tricarboxylic acid (TCA) cycle and aerobic respiration (Figure 2E). In contrast, genes whose expression is enriched in *Sntn*-negative lateral cells over *Sntn* cells include ribosomal genes, characteristic of more active cells (Llorens-Bobadilla et al., 2015; Shin et al., 2015). Indeed, a small fraction of captured lateral cells were proliferating, as inferred by the expression of cell cycle genes (e.g. *Mki67*) (Figure 2C and Table S4)*. Sntn*-negative lateral cells were also characterised by expression of *Bhmt*, a gene encoding for an enzyme involved in modulating gene expression through the BHMT-betain pathway and recently found to be key in the differentiation of immature oligodendrocytes (Sternbach et al., 2021). These subtle but noteworthy differences in the gene expression profiles of lateral ependymal cells suggest that these cells co-exist in immature and mature cell states in the adult mouse spinal cord.

Importantly, a large subset of immature *Sntn*-negative lateral cells expressed high levels of immediate early genes (including *Fos*, *Junb*, *Egr1*) and other stress-induced genes such as *Maff* and the growth factor gene *Ccn1* (Figure 2C and Table S4). Many of the genes enriched in this subset compared to the other *Sntn*-negative cells overlapped with a set of dissociation-induced genes in muscle stem cells (20 out of 31 dissociation-induced genes in muscle stem cells in common) (van den Brink et al., 2017), and thus it is likely that the tissue dissociation activated these lateral cells. Zeisel and colleagues noted a similar upregulation of immediate early genes in spinal cord ependymal cells (Zeisel et al., 2018). Biological processes overrepresented in these dissociation-affected cells included positive regulation of RNA metabolic process, regulation of cell cycle, and response to stress (Figure 2E and Table S4). Interestingly, some of these dissociation-induced genes were associated with spinal cord injury (Table S4), supporting the possibility that the dissociation indeed activated some immature lateral cells. Notably, despite their mature state, the transcriptomes of some *Sntn* cells were also affected by the tissue dissociation (Figure S3). Consistent with this, we found genes with higher expression in the mature cell cluster compared to the more immature cell cluster that were associated with response to endogenous stimulus, regulation of DNA-templated transcription in response to stress, response to cytokine, and cell cycle (Table S4). This raises the intriguing possibility that mature cells, despite their mature state, may be mobilised in response to spinal cord injury.

### The fraction of mature ependymal cells in the spinal cord increases with age

The ability of ependymal cells to proliferate and function as neural stem cells declines over time, which prompted us to investigate how ependymal cells change with age. To do so while minimising potential technical variability between samples, we generated a 10x scRNA-seq dataset from cells in and around the central canal region of aged wild-type mice (19-month-old mice, n = 4) in parallel to the young adult dataset discussed earlier (see Figure 1) (Figure 3A) (Methods). After aggregating the scRNA-seq libraries from young and aged mice and filtering out low-quality cells and doublets, we retained 19,354 cells (9,061 cells from young and 10,293 from aged mice). We recovered the same major cell types from the aged central canal as from the young central canal (Figure 3B); but observed substantial age-related heterogeneity within some of them (Figure S4A-C).

**Figure 3.**
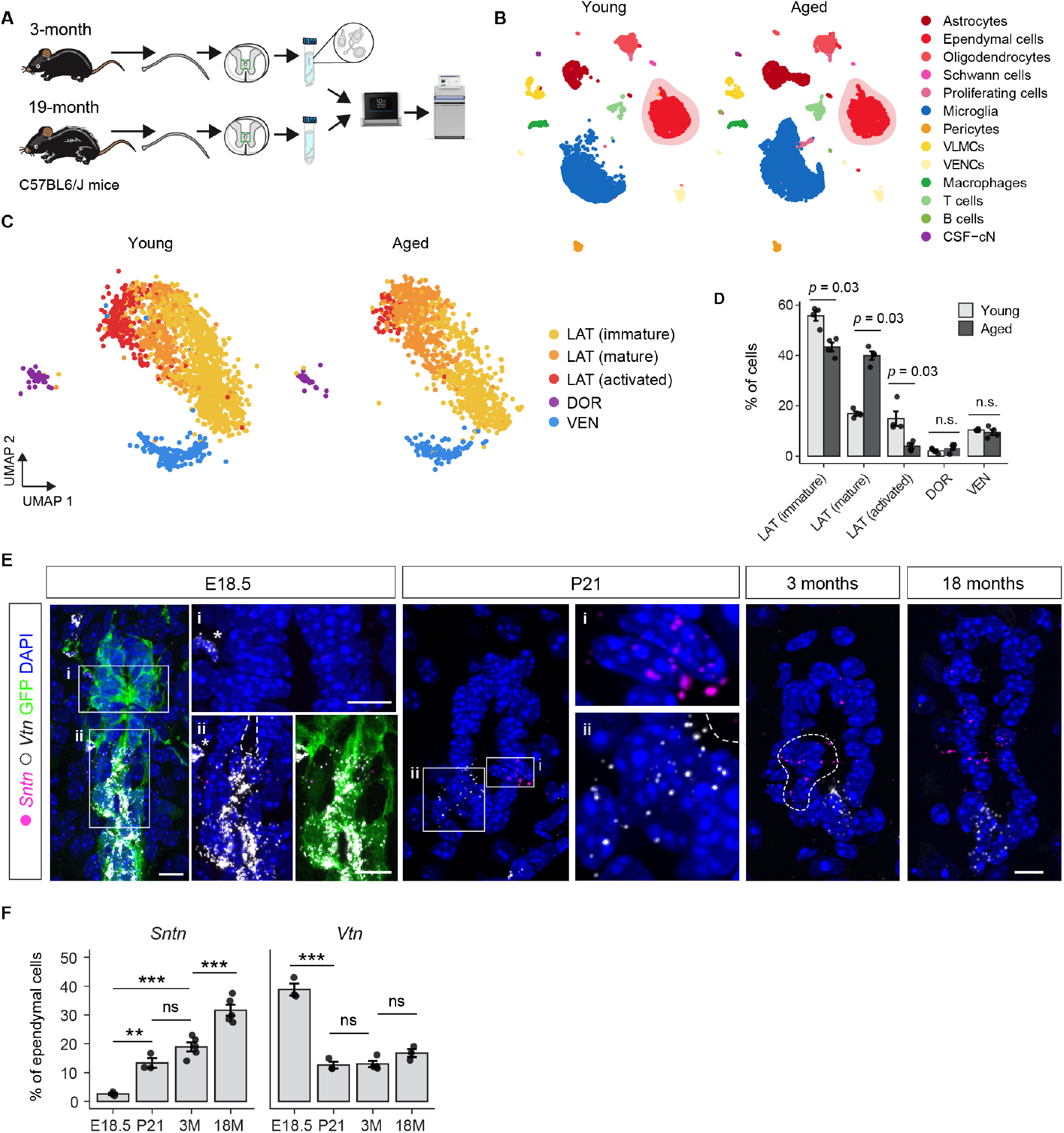
Age-related changes in spinal cord ependymal cells. (A) Workflow for the generation of the ageing dataset. (B) UMAP plot of spinal ependymal cells split by age and coloured by major cell types. (C) UMAP plot of ependymal cells subtypes and states from young and aged mice. (D) Bar plots showing the percentages of each ependymal cell subtype in young and aged mice. Each dot represents one mouse (n = 4). Bars indicate the mean, error bars standard deviation (SD), Wilcoxon rank sum test. (E) Images of the spinal cord central canal region showing cells expressing the mature lateral cell marker *Sntn* (magenta) and the ventral cell marker *Vtn* (grey) in E18.5 embryos, postnatal day (P) 21 pups, 3- and 18-month old mice. Images show a maximum intensity projection of 3-5 optical z-slices encompassing a cell length. The image from the E18.5 embryo additionally shows GFP expression under the control of the *Foxj1* promoter to better highlight ependymal cells. Note that a string of GFP^+^*Vtn*^+^ ventral cells extends from the central canal (high-magnification image in Fii), reaching all the way to the pia matter (not shown). Dashed line in image from a 3-month-old mouse encircles a typical cluster of *Sntn*-expressing cells. Scale bars: 20 μm and 5 μm (high-magnification images). (F) Quantification of the percentage of ependymal cells expressing *Sntn* and *Vtn* across ages. Each dot represents one mouse (n = 4). Bars indicate the mean, error bars standard deviation (SD), one-way ANOVA with Tukey’s multiple comparisons test (** p<0.01, *** p<0.001, ns is not significant).

To focus on age-related changes within the ependymal cell cluster, we extracted 3,527 ependymal cells (2,114 cells from young and 1,413 cells from aged mice), with a median of 8,285 UMIs and 3,119 genes detected per cell (Figure S4A). Clustering revealed that ependymal cell identity was largely maintained in the aged spinal cord (Figure 3C). Indeed, we identified all the ependymal cell subpopulations from 3-month-old mice in aged mice except the proliferating cell cluster, consistent with a decrease in ependymal cell proliferation with age (Alfaro-Cervello et al., 2012; Sabourin et al., 2009). To uncover potential age-related expression changes, we performed differential gene expression analysis between young and aged cells and found that ependymal cell transcriptomes from young and aged spinal cords were surprisingly similar. We only detected 54 age-regulated genes in cells from aged mice, 34 upregulated and 20 downregulated genes (Table S5). Genes whose expression was enriched in aged ependymal cells were associated with three main biological processes: mitochondrial function (e.g. *mt-Atp6*, *mt-Cytb*), which likely reflects the high energy requirements of fully functioning ependymal cells; glutathione metabolism (*Gstm1*, *Gpx3*, *Mgst1*), which plays important roles in antioxidant defence and nutrient metabolism and may be necessary for the proper functioning of metabolically active cells; and immune-related processes such as antigen processing and presentation and regulation of adaptive immune response (e.g. *Lyz2, Mif*), which suggests that ependymal cells play a bigger role than a purely mechanical barrier in protecting the CNS (see Table S5 for full list). Among the genes downregulated in aged cells were *Mt3*, a metallothionein protein involved in protecting against heavy metal toxicity; *Pcsk2*, a protease involved in the processing of hormone and other protein precursors; secreted protein genes *Mia* and *Ccn1*; and the cell cycle gene *Ccnd2* (cyclin D2). Most of the common age-regulated genes that we identified in spinal cord ependymal cells also change with age in brain ependymal cells (24/34 upregulated genes including *Caly*, *Lyz2*, and *Gstm1*, and 10/20 downregulated genes including *Ccnd2*, *Mia*, *Mt3*, *Pcsk2*) (Ximerakis et al., 2019).

Analysis of the relative abundance of each ependymal subtype in young and aged mice revealed that the percentage of the distinct lateral cell subpopulations shifted with age. Homeostatic and activated immature cells decreased from 56% to 43% and from 15% to 4%, while the percentage of mature cells increased from 17% in young mice to 40% in aged mice (Figure 3D). In contrast, the percentage of dorsal and ventral ependymal cells did not change postnatally (Figure 3D). To confirm and extend these observations, we investigated the expression of subtype-specific markers in the spinal cord across a wider range of ages using RNAscope (Figure 3E). Ependymal cells first appear in the developing mouse spinal cord between embryonic day (E) 15.5 and E17.5 and by E18.5 the ventricular layer of the developing spinal cord reaches the size of what will be the central canal (Canizares et al., 2019; Li et al., 2018). In the E18.5 spinal cord just ~3% of the ependymal cell population expressed *Sntn* (Figure 3E and F). The percentage of *Sntn*-expressing cells then continued to increase postnatally with 15% ependymal cells expressing *Sntn* at postnatal day (P) 21, 19% in 3-month-old mice, and 34% in 18-month-old mice (Figure 3E and F). Intriguingly, *Sntn*-expressing cells often appeared in clusters (Figure 3E). In comparison, the percentage of cells expressing the ventral-specific marker *Vtn* did not significantly change postnatally (Figure 3D and E). Importantly, RNAscope numbers are in line with scRNA-seq numbers indicating that our scRNA-seq dataset faithfully represents the cellular composition of the ependymal cell population and is not strongly affected by differences in the sensitivity of the cells to tissue dissociation or differences in cell capture. Taken together, our analysis identifies as the major age-related change within the spinal cord ependymal cell population the shift of lateral cells towards a *Sntn* phenotype, likely an indication of ongoing ependymal cell maturation in the adult mouse spinal cord.

### *Sntn* spinal cord ependymal cells are transcriptionally similar to postmitotic brain ependymal cells

While some ependymal cells in the spinal cord proliferate slowly or rarely postnatally and retain the ability to self-renew and give rise to specialised cell types in response to injury, brain ependymal cells are postmitotic and lack neural stem cell potential (Alfaro-Cervello et al., 2012; Carlén et al., 2009; Li et al., 2018; Meletis et al., 2008; Shah et al., 2018; Spassky et al., 2005; Stenudd et al., 2022). To uncover transcriptomic differences that may map to these important differences between spinal cord and brain ependymal cells, we integrated our 10x ependymal cell dataset from young mice with data from Zeisel and colleagues (Zeisel et al., 2018), which includes ependymal cell transcriptomes from across the CNS (Figure S5A and B) (Methods). The integrated dataset of 3,289 cells split into 15 clusters (Figure S5C and D) that we manually merged into 11 clusters based on their gene expression profile, known ependymal subtypes, and region of origin (Figure 4A and B).

**Figure 4.**
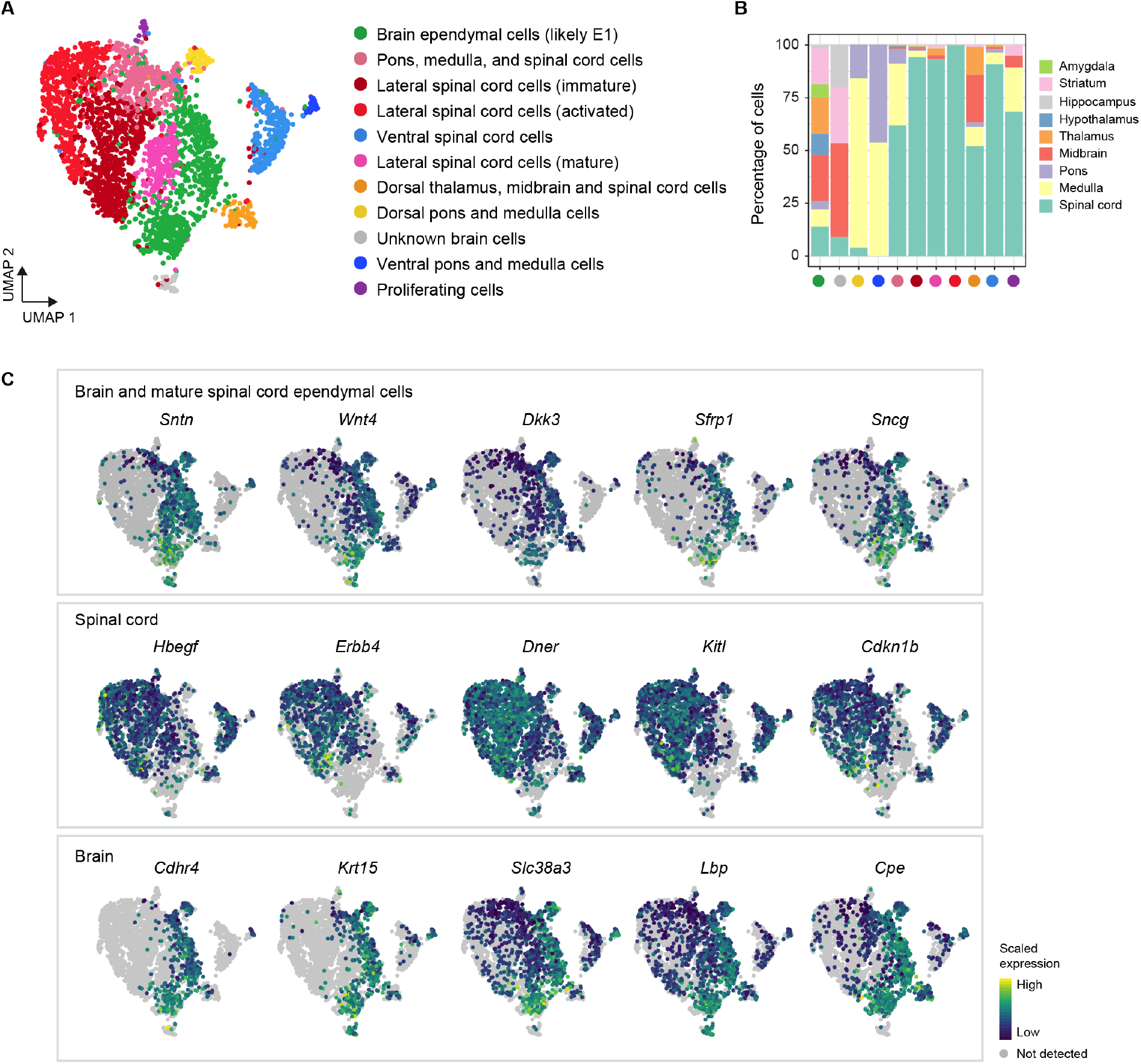
Ependymal cell heterogeneity across the CNS axis. (A) UMAP embedding of ependymal cells across the CNS, including cells from our spinal cord 10x dataset from young mice and cells from Zeisel et al. (Zeisel et al., 2018). (B) Stacked bar plot showing the percentage of cells within each cluster from each tissue. Dots colour-coded to represent ependymal subtype as shown in A. (C) UMAP plots showing the expression distribution of key genes.

Intriguingly, visualising how clusters split with increasing clustering resolution, we noticed that the mature spinal cord cell cluster arises from within the brain ependymal cell ‘branch’ (Figure S5C). This suggests that mature spinal cord cells are transcriptionally similar to brain ependymal cells. Indeed, *Sntn* and other genes expressed by mature ependymal cells in the spinal cord such as *Dkk3* and *Wnt4* were expressed by nearly all brain ependymal cells (Figure 4C, Table S6). The fact that ependymal cells in the mouse brain complete their maturation during the first two postnatal weeks (Spassky et al., 2005) further supports the idea that *Sntn* marks mature ependymal cells. Moreover, genes enriched in brain ependymal cells were associated with GO terms that were also overrepresented in mature spinal cord cells and in spinal cord ependymal cells from aged mice (e.g. mitochondrial function, cilia, and others suggesting a close interaction between brain ependymal cells and the immune system) (Table S6). These functional similarities between brain ependymal cells and *Sntn* spinal cord ependymal cells suggest that the latter represent a mature cell state and further support the observation that lateral spinal cord ependymal cells continue to mature throughout life although much more slowly than their brain counterparts.

Brain and spinal cord ependymal cells, including mature cells, also exhibited marked molecular differences (Figure 4C). Wnt/β-catenin signalling plays a key role in ependymal cell proliferation and tissue integrity in the spinal cord (Shinozuka et al., 2019; Xing et al., 2018). Intriguingly, brain ependymal cells uniquely expressed *Sfrp1*, a secreted molecule that blocks the interaction between Wnt ligands and their receptors and has recently been found to actively maintain quiescence in the subventricular zone (Donega et al., 2022) (Figure 4C). This identifies distinct regulation of Wnt signalling as a potential explanation for differences in maturation and proliferative capacity of brain and spinal cord ependymal cells. We also identified genes unique to spinal cord ependymal cells that indicate distinct signalling capabilities in the central canal niche. These include *Hbegf,* encoding a growth factor that acts via the epidermal growth factor receptor (EGFR) subfamily, and its potential receptor tyrosine kinase *Erbb4*; *Dner*, a paralog of *Notch4*; and *Kitl*, encoding the ligand for the receptor tyrosine kinase KIT (Figure 4C). Additionally, spinal cord ependymal cells uniquely express *Cdkn1b*, a cyclin-dependent kinase inhibitor involved in G1 arrest. In contrast, brain ependymal cells express specific genes related to cell adhesion (e.g. *Cdhr4, Dchs1*), cytoskeleton (e.g. *Krt15*, *Sncg*), transport (e.g. *Slc38a3*), innate immune response (e.g. *Lbp*), and hormone metabolism (e.g. *Cpe*). These comparisons provide further evidence for the protracted maturation of spinal cord ependymal cells and highlight maintenance of a more dynamic signalling microenvironment in the spinal cord. This raises the further intriguing possibility that the range of maturation states within the spinal cord central canal underlies the ability of this cell population to respond to injury, but that this ability is lost as cells mature.

### Mature ependymal cells are plastic and can re-enter the cell cycle in response to injury

To test whether mature spinal cord cells represent a terminally differentiated (i.e. postmitotic) cell state we next investigated how these cells respond to a challenge using an *ex vivo* model of spinal cord injury (Fernandez-Zafra et al., 2017). Briefly, we generated spinal cord slices from 5-month-old mice, and immediately fixed some slices to define the intact spinal cord conditions (day 0) while we cultured others for 3 and 5 days to investigate whether *Sntn* cells proliferate in this context (Figure 5A). To do so, we combined RNAscope for *Sntn* and the pan-ependymal marker *Mia* with immunofluorescence for the cell proliferation marker Ki67. In the intact spinal cord, 25% of all ependymal cells expressed *Sntn* and most appeared in clusters (Figure 5B and C). Of the few ependymal cells that we found proliferating in the intact spinal cord (3/862 or 0.4% of the ependymal cells) none expressed *Sntn* (Figure 5B and C). However, the number of Ki67 cells was so low that we cannot establish whether *Sntn* cells proliferate in the adult spinal cord under normal conditions. After 3 days, ependymal cell proliferation increased sharply in cultured spinal cord slices (40% of the cells) while the fraction of *Sntn*-expressing cells decreased significantly compared to normal conditions (from 25% to 14%) (Figure 5B and C). This suggested that either *Sntn*-expressing cells are preferentially lost following injury or that despite expression of this mature cell marker, such cells remain plastic and can downregulate *Sntn* in this context. In support of such cell plasticity, we found that the percentage of *Sntn* cells expressing low levels of *Sntn* (a single transcript/RNAscope dot) was higher in cultured slices than in the intact spinal cord (Figure 5C). Importantly, even though most Ki67-positive ependymal cells were *Sntn*-negative, a small fraction of low-*Sntn* cells were Ki67-positive (Figure 5B, day 3 inset i, arrowhead and quantified in 5C). Non-proliferating *Sntn* cells expressed high levels of *Sntn*, as they do in the intact spinal cord (Figure 5B, day 3 inset ii and compare with day 0 inset ii). These findings suggest that *Sntn* expression is dynamically regulated in some but perhaps not all mature lateral cells and that downregulation of *Sntn* (i.e. reversing maturation) is a pre-requisite to re-enter the cell cycle in this injury context.

**Figure 5.**
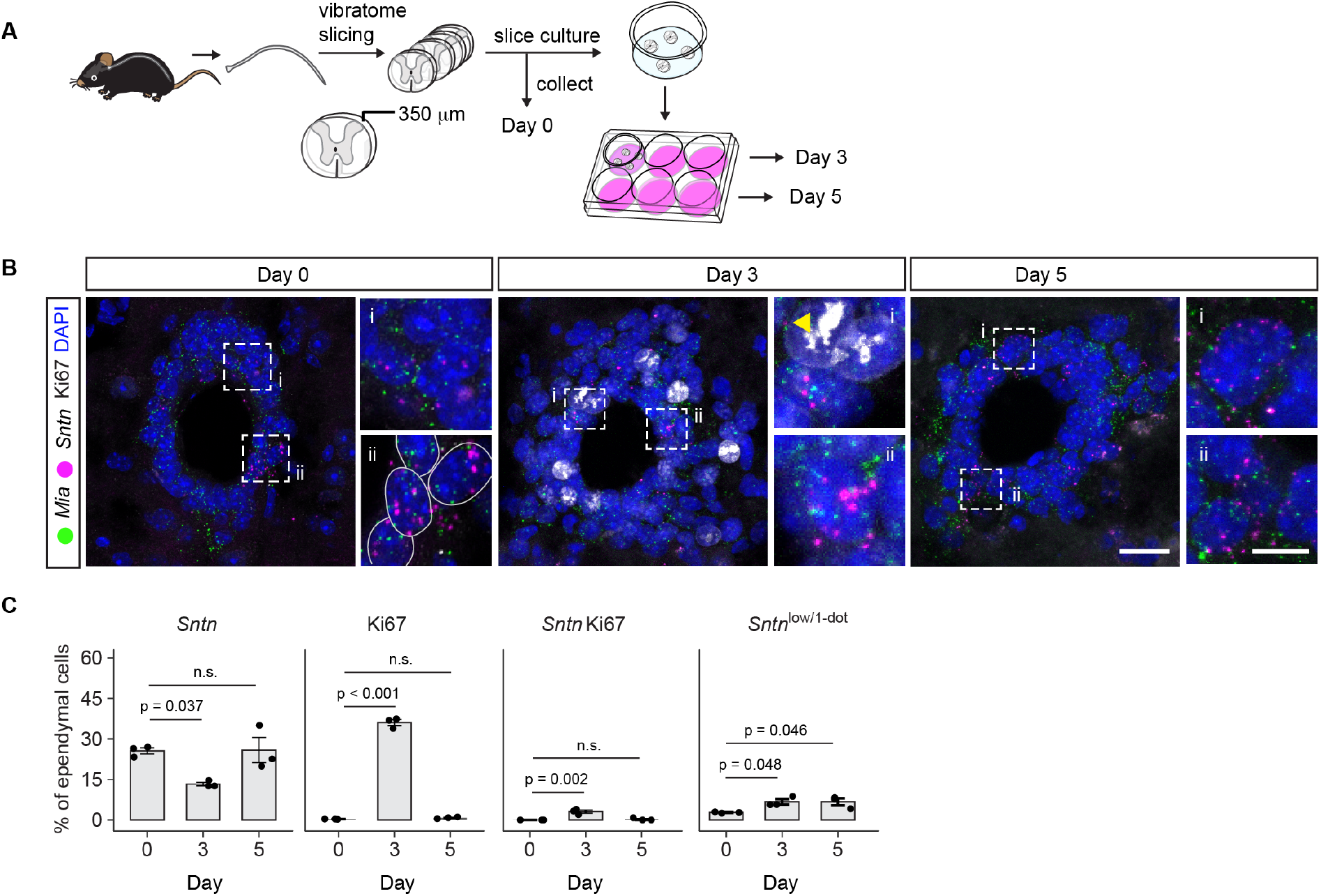
Mature ependymal cells downregulate *Sntn* and re-enter the cell cycle in response to injury. (A) Workflow of the spinal cord slice culture experiments. (B) Confocal images showing *Mia* and *Sntn* transcripts and Ki67 immunofluorescence on spinal cord slices fixed immediately after vibratome sectioning, at day 3 and day 5 of culture. Scale bar in the lower magnification images is 30 μm, in the higher magnification images is 10 μm. (C) Bar plots showing the percentage of *Sntn*-expressing ependymal cells, the percentage of Ki67 ependymal cells, the percentage of *Sntn* Ki67 cells, and the percentage of low *Sntn* cells (labelled with a RNAscope dot) day 0 (uninjured), day 3 and day 5 of culture. Each dot is a mouse and experiment (n = 3). A minimum of 201 cells were counted per mouse and time point. Bars indicate the mean, error bars standard deviation (SD), one-way ANOVA with Dunnett’s test (** p<0.01, *** p<0.001, ns is not significant).

Strikingly, in spinal cord slices cultured for 5 days the percentage of *Sntn*-expressing cells and proliferating cells were close to those under normal conditions (26% and 0.8% respectively). *Sntn* cells re-appeared in their characteristic clustered conformation (Figure 5B). Proliferating cells were again rare in slices cultured for 5 days (6/727 ependymal cells), but a few expressed low levels of *Sntn* (2/6 proliferating cells or 33%) (Figure 5B and C). None of the cells expressing high levels of *Sntn* were Ki67 positive. These findings do not rule out that mature lateral cells are quiescent under normal conditions but reveal that, in response to a challenge, *Sntn* expression is dynamically regulated in at least some mature cells and that these can then re-enter the cell cycle. The finding that the proportion of *Sntn* cells around the central canal was restored soon after the burst of injury-induced proliferation and that the integrity of the canal was preserved throughout the culture highlights the regulative properties of the central canal niche.

### Ependymal cells in the adult human spinal cord express SNTN and become ventralised with age

To determine the relevance of our findings in the mouse to the human spinal cord, we next studied *SNTN* expression in human samples ranging from the third to the sixth decade of life (years 20-29, 30-39, 40-49, 50-59, a minimum of three samples for each age group) (Table S7). We used FOXJ1 immunofluorescence to identify ependymal cells and RNAscope to detect expression of *SNTN* and of the broadly expressed gene *PPIB* to control for RNA quality and non-specific probe binding (as overlapping *SNTN* and *PPIB* signal) (Figure 6A). Across all ages, we found a disorganised ependyma and varying degrees of central canal stenosis in the human spinal cord, with more samples from young adults (years 20-29) having a patent canal (3/4 samples) (Figure 6, Table S7). *SNTN* was widely expressed in all the samples examined (Figure 6A). In contrast to the mouse, the fraction of *SNTN*-expressing ependymal cells appeared higher in samples from young adults (20-29 years) and to progressively decline in samples from older age groups (Figure 6). Quantification of the fraction of *SNTN* cells, however, was challenging due to tight packing of ependymal cells, differing sample quality, and autofluorescence, especially in tissue from older age groups.

**Figure 6.**
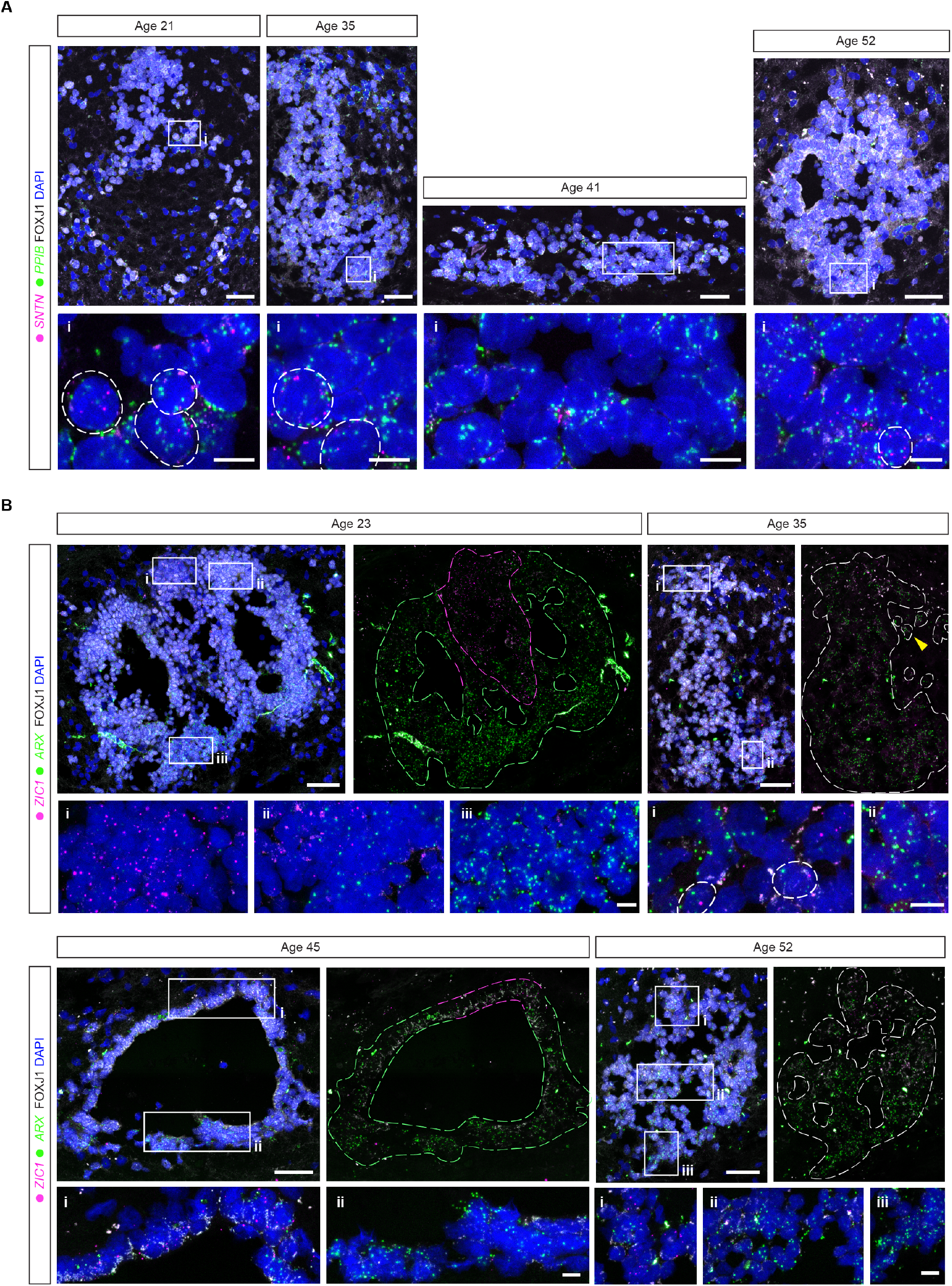
Ependymal cell identities in the human spinal cord. (A) Confocal images of the ependymal region from the human spinal cord of each age group, showing combined *SNTN* (magenta) and *PPIB* (green) RNAscope and FOXJ1 immunofluorescence. Insets below show the close-up of the regions indicated in their respective lower magnification images. The circles in the insets delineate ependymal cell nuclei. Scale bars in the lower magnification images are 50 μm, in the higher magnification images is 10 μm. (B) Confocal images of the ependymal region from the human spinal cord of each age group, showing combined RNAscope against *ZIC1* (magenta) and *ARX* (green) and FOXJ1 immunofluorescence and RNAscope signal only (right panel within each age sample). The dashed lines delineate the ependymal region to better appreciate the distinct domains. Arrowhead in the sample age 35 points to a few ependymal cells expressing *ARX* that have detached from the main cluster of ependymal cells as reported earlier (Alfaro-Cervello et al., 2014; Dromard et al., 2008). Insets show the close-up of the regions indicated in their respective lower magnification images. Scale bars in the lower magnification images are 50 μm, in the higher magnification images is 10 μm. All the images are maximum intensity projections of 5-6 1 μm optical sections, or about one cell length. Details about the samples analysed are summarised in Table S7.

Our mouse data indicated that dorsal and ventral ependymal cells continue to act as signalling centres in the adult spinal cord and their position may also uncover if and how the central canal niche microenvironment and cellular composition change with age. We therefore also performed RNAscope assays to detect transcripts for dorsal *ZIC1*-expressing and ventral *ARX*-expressing ependymal cells. Surprisingly, we found near complementary *ZIC1* and *ARX* domains across all ages examined, with most ependymal cells expressing *ARX* and a smaller subset of cells expressing *ZIC1* at the dorsal pole (Figure 6B). This was independent of the level of central canal stenosis and, even in samples with a patent canal, *ARX*-expressing ependymal cells abutted dorsal *ZIC1*-expressing cells (see Figure 6B, age 45). At the boundary between *ARX* and *ZIC1* domains we often found ependymal cells co-expressing *ARX* and *ZIC1* (Figure 6B, inset from sample age 35). As noted above, quantification of the fraction of cells expressing each marker was challenging, however, from age 30 fewer cells appeared to express *ZIC1* and did so at lower levels. In contrast, *ARX* was expressed consistently by most ependymal cells, with cells located ventrally expressing the highest levels. Together, these findings indicate that, unlike in the mouse, ependymal cells in the human spinal cord become ventralised with age.

## Discussion

In this study, we comprehensively characterise ependymal cell heterogeneity in the adult mouse spinal cord and relate this to the analogous human cell population. Our data divide mouse ependymal cells into dorsal, ventral, and lateral cell subtypes based on patterning genes that reflect embryonic origins and define functions. We find that many lateral cells mature slowly, in contrast to their brain counterparts, linking the range of cell maturation states to the ability of ependymal cells to respond to injury. Using an *ex vivo* model of spinal cord injury, we discover that lateral cell maturation is reversible and short-lived, with cells rapidly re-establishing a mature cell state. In striking contrast to the mouse, we report that almost all human spinal cord ependymal cells express a mature cell marker and acquire a ventral identity with age. This study provides a new resource with which to uncover the potential of different ependymal cell subtypes to drive spinal cord repair, it circumscribes the cell population that harbours the elusive neural stem cell potential and highlights the regulation of cell maturity as a key mechanism that provisions and perhaps limits response to injury in mammals.

A core finding from our study is that lateral ependymal cells in the mouse spinal cord uniquely exist in immature and mature cells states. Indeed, we found that the lateral cell population shifted towards a mature state over time, but immature cells remained a significant fraction of the central canal even in aged mice. This contrasts with the more rapid maturation of brain ependymal cells, which in the mouse is complete two weeks after birth (Spassky et al., 2005). Importantly, while mature spinal cord cells and brain ependymal cells share defining molecular features (Figure 5), it is clear that some spinal cord ependymal cells uniquely behave as neural stem cells (Barnabé-Heider et al., 2010; Meletis et al., 2008; Shah et al., 2018; Stenudd et al., 2022). Our findings therefore raised the possibility that this elusive neural stem potential resides within the immature lateral cell fraction and is lost as cells fully mature. We tested this hypothesis in an *ex vivo* model of spinal cord injury and found that the mature spinal cord ependymal cell state is plastic, with mature cells responding to injury and adapting to reinstate tissue homeostasis. Consistent with this, spinal cord injury triggers the re-coupling of ependymal cells in a way that resembles neonatal connectivity (Fabbiani et al., 2020). Moreover, we found that cell identity was highly regulated in the central canal as the fraction of mature cells returned to normal levels within a few days. These *ex vivo* findings align well with those in a recent report comparing scRNA-seq datasets for whole uninjured and injured mouse spinal cord (Milich et al., 2021). This study identified two ependymal cell subtypes, Ependymal-A and Ependymal-B, that correspond to our mature and immature lateral cells respectively, and also found that cilia-related gene expression and the fraction of Ependymal-A/mature cells transiently decreases following injury (Milich et al., 2021) (see also (Chevreau et al., 2021)).

As we were preparing this paper for publication further scRNA-seq evidence for ependymal cell downregulation of cilia-related genes after injury was reported (Stenudd et al., 2022). This study focused on *Tnfrsf19* (Troy)-expressing ependymal cells, some of which displayed extraordinary neural stem cell capabilities *in vitro*. However, consistent with RNAscope data in Stenudd et al., 2022, we found that cells of every ependymal subtype and state identified in our comprehensive scRNA-seq dataset express *Tnfrsf19* (Figure S6). This indicates that Troy-CreER recombined cells include ependymal cells with distinct identities, and so it remains unclear which of these possess neural stem cell properties. On the other hand, some clues are provided by the distinct behaviours of Troy-CreER recombined cells after injury: clones with one or more ependymal cells and one or more migrating/differentiating daughter cell/s were located dorsally or laterally, while others formed small ependymal cell clusters without obvious migrating progeny. The lack of migrating progeny was also characteristic of ventral recombined cells (Stenudd et al., 2022). This indicates some lateral and ventral ependymal cells lack the characteristic stem cell clonal pattern (a resident ependymal cell and a migrating/differentiating progeny, indicative of self-renewal and lineage/differentiation potential). Moreover, the clones of clustered ependymal cells are reminiscent of the clusters of mature lateral cells that we found quickly re-appearing after the burst in injury-induced cell proliferation (Figure 5C) and may indicate that the neural stem cell potential of mature lateral cells is limited.

As we find that many lateral ependymal cells are immature in the adult spinal cord these are likely to be the cells that mount a stem cell-like response to injury, as most Ki67-positive cells lack *Sntn* expression in our *ex vivo* assay. However, the downregulation of *Sntn* in some mature cells clearly also links regulation of cell maturity to this response. In unchallenged conditions mature cells are also localised in clusters (Figure 2D) and, in addition to *Sntn*, express the secreted Wnt antagonist *Dkk3* and the non-canonical Wnt ligand *Wnt4* and are often close to a *Dkk3*-expressing neuron (data not shown). In the spinal cord, canonical Wnt/β-catenin signalling regulates ependymal cell proliferation and maintains integrity of the ependymal epithelium (Shinozuka et al., 2019; Xing et al., 2018). These findings therefore suggest that mature cells locally inhibit Wnt/β-catenin signalling and induce quiescence in neighbouring cells, while their dedifferentiation may derepress Wnt/β-catenin and liberate neural stem cell behaviour. This scenario may explain how neural stem cell potential is eventually lost with increasing mature cell numbers and their deeper quiescence that come with age; such local inhibition in vivo may also explain the greater neural stem cell abilities that spinal cord ependymal cells display when cultured *in vitro*. Importantly, similar mechanisms operate in various stem cell niches: in the brain subventricular zone, *Dkk3* inhibits Wnt/β-catenin signalling (Zhu et al., 2014) and WNT4 contributes to regulation of neural stem cell quiescence (Chavali et al., 2018); WNT4 secreted by myofibers also induces muscle stem cell quiescence (Eliazer et al., 2019; Otto et al., 2008). Investigating the fine-tuning of Wnt signalling in the central canal niche will be key for understanding how the balance between self-renewal and quiescence is preserved or re-gained among ependymal cells. When this balance is lost, uncontrolled ependymal cell proliferation can lead to ependymal tumours also known as ependymomas, which notably include cells with a marker gene profile similar to that of activated lateral cells (Gillen et al., 2020; Gojo et al., 2020). The activation of lateral cells in our dataset was triggered by experimental procedure, but the decreasing fraction of these cells with age is intriguing given that ependymomas are often childhood cancers. Understanding how ependymal cell proliferation is kept in check may therefore inform cancer treatments as well as directed regenerative therapies.

How mouse ependymal cells relate to those in humans is poorly understood. Obtaining high-quality human spinal cord samples is obviously challenging and a good number are needed given the variable morphological changes that take place with age (Alfaro-Cervello et al., 2014; Garcia-Ovejero et al., 2015; Kasantikul et al., 1979; Milhorat et al., 1994; Yasui et al., 1999). Ependymal cells in the adult human spinal cord appear to retain expression of dorsoventral patterning genes as in the mouse, however this was observed in just one, teenage, sample (Ghazale et al., 2019). Based on *SNTN* expression, we show that most ependymal cells in the human spinal cord from age 20 are in a mature state (Figure 6A). This correlates with the apparent lack of ependymal cell proliferation in the adult human spinal cord (Alfaro-Cervello et al., 2014; Dromard et al., 2008). Unexpectedly, we found that the ventral marker *ARX* and the dorsal marker *ZIC1* are expressed in abutting domains in the adult human central canal, with virtually all cells located ventrally and laterally expressing *ARX* and a much smaller fraction of dorsally located cells expressing *ZIC1*. This contrasts strikingly with expression patterns in the mouse, where *Arx* and *Zic1* domains do not change throughout life (Figure 3) and suggests that human ependymal cells become ventralised with age. Intriguingly, such ventralisation resembles that of lizard spinal cord ependymal cells, which do not retain dorsoventral patterning genes and instead display a ventral identity (Lozito et al., 2021; Sun et al., 2018). This leads to imperfect regeneration when lizards lose their tails, as the spinal cord and surrounding tissues regrow with a ventral character due to the restricted potential of lizard ependymal cells and the signalling microenvironment they create (Lozito et al., 2021; Sun et al., 2018). Our findings raise intriguing questions about how the ventralisation of ependymal cells changes their potential and signalling microenvironment in the human spinal cord, and how this may impact cell transplantation strategies. In the mouse, an elegant study found chromatin accessibility for a latent oligodendrocyte differentiation programme in at least some ependymal cells and demonstrated how these cells can be nudged towards contributing oligodendrocytes to the injured spinal cord and promote repair (Llorens-Bobadilla et al., 2020). In the light of our findings in mouse and humans, it will be important to investigate whether that window of opportunity is lost with cell maturation and if manipulation of distinct ependymal cell subtypes can be used to therapeutic effect.

## Methods

No statistical methods were used to predetermine sample size. The experiments were not randomised, and investigators were not blinded to allocation during experiments and outcome assessment.

### Resource availability

#### Lead contact

Further information and requests for resources and reagents should be directed to the lead contacts, Aida Rodrigo Albors and Kate G. Storey.

#### Materials availability

This study did not generate new unique reagents.

### Experimental model and subject details

#### Mice

FOXJ1-EGFP transgenic mice (Ostrowski et al., 2003) were used for the generation of the Smart-seq2 scRNA-seq dataset (3-month-old) and for RNAscope experiments (E18.5, P21, and 3-month-old). For timed matings, the morning of the plug was considered E0.5. C57BL/6J wild-type mice (Charles River) were used for the generation of 10x scRNA-seq datasets (3- and 19-month-old mice), for RNAscope (18-month-old mice) and spinal cord slice culture experiments (5-month-old mice). All mice used in scRNA-seq experiments were females, while female and male mice were used for RNAscope and slice culture experiments. Animals were housed in standard housing conditions on a 14 h light/10 h dark cycle with food and water ad libitum. All animal procedures were performed in accordance with UK and EU legislation and guidance on animal use in bioscience research. This work was performed under the UK project license 60/4454 and P0B7F2E8D and subjected to local ethical review.

#### Human samples

The spinal cord samples were obtained from The Netherlands Brain Bank, Netherlands Institute for Neuroscience, Amsterdam (open access: http://www.brainbank.nl/). All material has been collected from donors for or from whom a written informed consent for a brain autopsy and the use of the material and clinical information for research purposes had been obtained by the NBB. An overview of the clinical information and post-mortem variables of donors in this study is summarised in Table S7.

### Method details

#### Cell preparation for scRNA-seq

##### Cell preparation for plate-based scRNA-seq (Smart-seq2)

For the plate-based scRNA-seq dataset, we generated cell preparations from 3-month-old FOXJ1-EGFP mice. A total of four cell preparations were generated, one per day and starting at 10 AM. In parallel to the FOXJ1-EGFP cell preparation we generated a cell preparation from C57BL/6J wild-type mice to set the GFP threshold for sorting. Mice were deeply anaesthetised with an intraperitoneal injection of pentobarbital before being transcardially perfused with ice-cold artificial cerebrospinal fluid (aCSF) solution (87 mM NaCl, 2.5 mM KCl, 1.25 mM NaH2PO4, 26 mM NaHCO3, 25 mM glucose, 1 mM CaCl2, 2 mM MgSO4, 10 mM HEPES in sterile MilliQ water, pH 7.4) that was oxygenated in 95% O2 5% CO2 until used. After laminectomy, the spinal cord was carefully dissected out and transferred to a Sylgard-coated dish filled with ice-cold aCSF for microdissection. The thoracic region of the spinal cord was separated with a scalpel and pinned, ventral side down, then cut through the midline in two halves as in an open-book preparation. Working under a fluorescence stereo microscope, the central canal of the spinal cord was identified as an opaque stripe running along the middle of either of the halves of the spinal cord or, aided by fluorescence, as a bright GFP stripe. The central canal was quickly microdissected, cut in small pieces, and placed in aCSF on ice until the whole thoracic central canal was microdissected. Quickly, the aCSF was replaced with 1 mL of pre-warmed digestion solution (3X TrypLE Select enzyme (Thermo Fisher) diluted in Hanks’ Balanced Salt Solution (HBSS) without calcium and magnesium, 15 mM HEPES, and 10 μg/ml DNase I (Worthington)) and the tissue pieces incubated in a water bath at 37 ºC. After 45 min, tissue pieces were gently pipetted up and down 10 times to facilitate their dissociation and immediately returned to the water bath. After 20 min, the remaining tissue pieces and cells were gently pipetted up and down 12x with a P1000 pipette tip and 22x with a P200 pipette tip. The enzymatic digestion was then stopped with wash solution (HBSS, 15 mM HEPES, 0.05% BSA, 0.5 mM EDTA) and the cell suspension filtered through a 50 µm cell strainer (CellTrics, Sysmex Europe) and centrifuged at 300 xg for 5 min. After discarding the supernatant, myelin debris was removed from the cell suspension using Myelin Removal Beads II kit (Miltenyi Biotec) following the manufacturer’s protocol with minor modifications. Briefly, the cell suspension was incubated with the adequate volume of Myelin Removal Beads II for 15 min in the fridge (4 ºC), then washed well by adding 10x the beads volume of MACS buffer and centrifuged at 300 xg for 5 min. LS columns with a 30 µm pre-separation filter (Miltenyi Biotec) were equilibrated with MACS buffer on a QuadroMACS magnetic separator (Miltenyi Biotec) and as soon as the cell suspension finished centrifugating, cells were passed through the filter and column to remove cell clumps and myelin debris. The cell suspension was then centrifuged at 300 xg for 5 min and the MACS buffer quickly replaced with sorting buffer (HBSS, 15 mM HEPES, 0.5% BSA, 1 mM EDTA, 5 µg/mL 4,6-diamidino-2-phenylindole (DAPI)) and the cell pellet gently resuspended and placed on ice until cells were sorted into lysis buffer plates. Lysis plates were created by dispensing 0.4 μl lysis buffer (0.5 U Recombinant RNase Inhibitor (Takara Bio, 2313B), 0.0625% Triton X-100 (Sigma, 93443-100ML), 3.125 mM dNTP mix (Thermo Fisher, R0193), 3.125 μM Oligo-dT 30 VN (IDT, 5′AAGCAGTGGTATCAACGCAGAGTACT 30 VN-3′) and 1:600,000 ERCC RNA spike-in mix (Thermo Fisher, 4456740)) into 384-well hard-shell PCR plates (Biorad HSP3901) using a Tempest liquid handler (Formulatrix). All plates were then spun down for 1 min at 3220 xg and snap frozen on dry ice. Plates were stored at −80°C until used for sorting.

GFP-positive DAPI-negative cells were sorted into lysis plates with a BD Influx Cell Sorter using 6 psi pressure and a 130 μm nozzle. Flow cytometry data was analysed using BD FACSDiva software. Sorted plates were immediately spun down for 1 min and snap frozen on dry ice until processed. To maximise cell recovery, wide-mouth low-retention tips pipette tips were used through the cell preparation. Low retention microcentrifuge/Eppendorf tubes were also used throughout. All working solutions were adjusted to pH 7.4. and kept on ice, except the digestion solution that was kept at 37 ºC. Spinal cord tissue and cells were kept on ice at all times except during enzymatic digestion.

##### Cell preparation for droplet-based scRNA-seq (10x Genomics)

For the droplet-based scRNA-seq datasets, we generated samples from 4 young (3-month-old) and 4 aged (19-month-old) C57BL/6J mice, with two animals killed per day: one young and one old. To avoid introducing technical bias (Baran-Gale et al., 2018), the two cell preparations were processed in parallel, alternating which sample was processed first (alternating between 3-month old sample followed by the 19-month old sample, and then the 19-month sample followed by the 3-month sample). The cell preparation for 10x scRNA-seq was based on that for plate-based scRNA-seq, with some modifications to improve the cell yield and minimise cell clumps: (1) all the solutions used for cell preparations for 10x scRNA-seq contained 1% (w/v) trehalose as we found that it greatly improved cell survival (Saxena et al., 2012) and thus cell yield, (2) we added 50 µg/mL (0.28 collagenase Wünsch units) of Liberase TM (Merck) in the digestion solution, and (3) added an additional digestion step after the myelin removal step. We also added a step to lyse red blood cells with the goal of maximising ependymal cell capture. Briefly, mice were deeply anaesthetised and perfused transcardially with ice-cold, oxygenated aCSF. The spinal cords were carefully dissected out and transferred onto a dish filled with ice-cold aCSF for microdissection. For 10x scRNA-seq cell preparations the whole spinal cord was used, and the central canal was identified as an opaque stripe running along the middle of the spinal cord in an open-book preparation. Microdissected central canal tissue pieces were kept in aCSF on ice until the whole or most of the spinal cord central canal was microdissected for both samples. Tissue pieces were then incubated in pre-warmed digestion solution (3X TrypLE Select enzyme and 50 µg/mL Liberase TM diluted in HBSS without calcium and magnesium, 15 mM HEPES, and 10 µg/ml DNase I, 1% (w/v) trehalose) and incubated in an incubator shaker at 37 ºC. Every 15 min, tissue pieces were gently pipetted up and down 10x to facilitate their dissociation. After 65 min of digestion, cell preparations were gently pipetted up and down 12x with a P1000 pipette tip and 22x with a P200 pipette tip and wash solution (HBSS without calcium and magnesium, 15 mM HEPES, 0.4% BSA, 0.1% trehalose) was added to stop the digestions. Cell suspensions were then filtered through a 50 µm cell strainer (CellTrics, Sysmex Europe) and centrifuged at 300 xg for 6 min at 4 ºC. Cell pellets were resuspended in red blood cell lysis buffer (Roche, cat.# 11814389001) and incubated for 1 min at RT. MACS buffer was then added and cell suspensions centrifuged at 300 xg for 6 min. Myelin debris was removed from the cell suspensions using Myelin Removal Beads II kit (Miltenyi Biotec) as described above but with 1 min longer centrifugation steps. Note that this step seems to deplete most neurons from the cell preparation and thus neurons are underrepresented in 10x datasets. It is also likely that neurons do not survive well our dissociation protocol. Following myelin removal, cell suspensions were incubated in post-digestion solution (3X TrypLE Select enzyme, HBSS without calcium and magnesium, 15mM HEPES, 1 mM EDTA, 1% (w/v) trehalose) for 8 min in the 37 ºC incubator shaker. The digestions were stopped with wash solution without EDTA and immediately centrifuged at 300 xg for 4 min. Cell pellets were quickly but gently resuspended in wash buffer without EDTA and kept on ice until loaded into the Chromium controller (10x Genomics). A small aliquot of each cell suspension was mixed with the same volume of Trypan Blue Solution (0.4%) (Thermo Fisher) to estimate cell concentration and cell viability before loading the cell preparations into the Chromium chip (10x Genomics).

#### Single-cell library preparation and sequencing

##### Plate-based scRNA-seq

cDNA synthesis was performed using the Smart-seq2 protocol (Picelli et al., 2013, 2014). Briefly, 384-well plates containing single-cell lysates were thawed on ice followed by first strand synthesis. 0.6 μl of reaction mix (16.7 U/μl SMARTScribe TM Reverse Transcriptase (Takara Bio, 639538), 1.67 U/μl Recombinant RNase Inhibitor (Takara Bio, 2313B), 1.67X First-Strand Buffer (Takara Bio, 639538), 1.67 μM TSO (Exiqon, 5′-AAGCAGTGGTATCAACGCAGACTACATrGrG+G-3′), 8.33 mM DTT (Bioworld, 40420001-1), 1.67 M Betaine (Sigma, B0300-5VL), and 10 mM MgCl 2 (Sigma, M1028-10X1ML)) was added to each well using a Tempest liquid handler. Bulk wells received twice the amount of RT mix (1.2 μl). Reverse transcription was carried out by incubating wells on a ProFlex 2×384 thermal-cycler (Thermo Fisher) at 42°C for 90 min and stopped by heating at 70°C for 5 min. Subsequently, 1.6 μl of PCR mix (1.67X KAPA HiFi HotStart ReadyMix (Kapa Biosystems, KK2602), 0.17 μM IS PCR primer (IDT, 5′-AAGCAGTGGTATCAACGCAGAGT-3′), and 0.038U/μl Lambda Exonuclease (NEB, M0262L)) was added to each well with a Tempest liquid handler (Formulatrix). The amplified product was diluted with a ratio of 1 part cDNA to 10 parts 10mM Tris-HCl (Thermo Fisher, 15568025), and concentrations were measured with dye-fluorescence assay (Quant-iT dsDNA High Sensitivity kit; Thermo Fisher, Q33120) on a SpectraMax i3x microplate reader (Molecular Devices). These wells were reformatted to a new 384-well plate at a concentration of 0.3 ng/μl and final volume of 0.4 μl using an Echo 550 acoustic liquid dispenser (Labcyte). If the cell concentration was below 0.3 ng/μl, 0.4 μl of sample was transferred.

Illumina sequencing libraries were prepared using the Nextera XT Library Sample Preparation kit (Illumina, FC-131-1096) (Darmanis et al., 2015; Schaum et al., 2018). Each well was mixed with 0.8 μl Nextera tagmentation DNA buffer (Illumina) and 0.4 μl Tn5 enzyme (Illumina), then tagmented at 55°C for 10 min. The reaction was stopped by adding 0.4 μl “Neutralize Tagment Buffer” (Illumina) and spinning at room temperature (RT) in a centrifuge at 3220 xg for 5 min. Indexing PCR reactions were performed by adding 0.4 μl of 5 μM i5 indexing primer, 0.4 μl of 5 μM i7 indexing primer, and 1.2 μl of Nextera NPM mix (Illumina). PCR amplification was carried out on a ProFlex 2×384 thermal cycler using the following program: 1. 72°C for 3 minutes, 2. 95°C for 30 s, 3. 12 cycles of 95°C for 10 s, 55°C for 30 s, and 72°C for 1 min, and 4. 72°C for 5 min.

Following library preparation, wells of each library plate were pooled using a Mosquito liquid handler (TTP Labtech). Pooling was followed by two purifications using 0.7x AMPure beads (Fisher, A63881). Library quality was assessed using capillary electrophoresis on a Fragment Analyzer (AATI), and libraries were quantified by qPCR (Kapa Biosystems, KK4923) on a CFX96 Touch Real-Time PCR Detection System (Biorad). Plate pools were normalized to 2 nM and sequenced on the NovaSeq 6000 Sequencing System (Illumina) using 2×100bp paired-end reads with an S4 300 cycle kit (Illumina, 20012866).

##### Droplet-based scRNA-seq (10x Genomics)

cDNA libraries were generated with 10x Genomics Chromium Single Cell 3′ Library and Gel Bead Kit v3.1 following the manufacturer’s instructions. Quality control of the cDNA and sequencing libraries was performed using a TapeStation System (Agilent) with High Sensitivity D5000 ScreenTape and library concentrations were measured with Qubit dsDNA High Sensitivity assay kit (Thermo Fisher Scientific). The eight libraries (one for each mouse) were then pooled for sequencing, taking into account the expected differences in cell numbers between each library. Sequencing was peformed by Edinburgh Genomics (University of Edinburgh) on one lane of a NovaSeq 6000 Sequencing System (Illumina) with the following cycle setup for paired-end reads: read 1, 28 cycles; i7 index, 8 cycles; read 2, 91 cycles).

#### scRNA-seq data preprocessing

##### Plate-based scRNA-seq (Smart-seq2)

Raw sequencing reads were trimmed to remove contaminating adapter sequences and low-quality ends (using a cutoff threshold of 15) with cutadapt v1.16 (Martin, 2011). Reads shorter than 75 nucleotides were filtered out. Trimmed and paired reads were then aligned to the mouse reference genome (mm10, Ensembl release 98) enhanced with ERCC spike-in and EGFP sequences and quantified using the STARsolo pipeline within STAR 2.7.7a (Dobin et al., 2013; Kaminow et al., 2021). The output of STARsolo is similar to that of Cell Ranger (Kaminow et al., 2021), which we used to generate the count matrices from the 10x data. STAR parameters were: --soloType SmartSeq --soloUMIdedup Exact --soloStrand Unstranded --sjdbOverhang 99 --outFilterType BySJout --outFilterMultimapNmax 20 --outSAMtype BAM SortedByCoordinate --outSAMattributes All --outSAMunmapped Within KeepPairs --quantMode GeneCounts TranscriptomeSAM. The rest of parameters were set to default values. Gene count matrices were then loaded into Seurat v3.2.2 (Stuart et al., 2019) for further analysis.

##### Droplet-based scRNA-seq (10x)

Raw sequencing reads were preprocessed with Cell Ranger v4.0 (10x Genomics) following the manufacturer’s instructions with default parameters and aligned to the mouse reference genome (mm10, Ensembl release 100). The output filtered feature-barcoded matrices (count matrices) were then loaded into Seurat v3.2.2 (Stuart et al., 2019) for further analysis.

#### scRNA-seq quality control, normalisation, and dimensionality reduction

All gene count matrices were merged by dataset and processed using Seurat v3.2.2 (Stuart et al., 2019). Cells expressing fewer than 500 genes and genes expressed in fewer than 3 cells were removed. Smart-seq2 data were normalised with a scale factor of 1,000,000 and 10x data were normalised with a scale factor 10,000 then log transformed using Seurat’s NormalizeData function. Highly variable features were identified with the FindVariableFeatures function using the vst method with default parameters. After scaling and centring the data, the top 2,500 highly variable features were used for principal component analysis (PCA) and significant principal components (PCs) were used for graph-based clustering (shared nearest neighbour graph and Louvain clustering using Seurat’s FindClusters function). The same PCs were used as input to uniform manifold approximation and projection (UMAP) (McInnes et al., 2018) for data visualisation. Clustering was performed at a range of clustering resolutions and the R package clustree (Zappia and Oshlack, 2018) was used to visualise and understand relationships between clusters, in combination with differential expression analyses. The most biologically meaningful clustering resolution was chosen. To annotate cell clusters, differential expression analyses were performed using a Wilcoxon rank sum test. Cell clusters were manually annotated based on known marker genes and data from http://www.mousebrain.org (Zeisel et al., 2018). Clusters that co-expressed mutually exclusive markers (i.e. markers that define distinct cell types) were considered doublets and removed from the datasets. A few, small non-ependymal cell clusters (e.g. microglia) were removed from the Smartseq2 dataset. After removing potential doublets and contaminating cells, highly variable features, PCA, clustering and visualisation was repeated as described above until no more doublets or contaminating cells remained in the datasets. The resulting datasets were aggregated or integrated as described below.

#### Ependymal cell heterogeneity analysis

After initial quality control, filtering, and annotating the 10x dataset of cells from the spinal cord central canal region of young adult mice, we subset the ependymal cell cluster to integrate it with the Smart-seq2 ependymal cell dataset using Seurat’s CCA (Stuart et al., 2019). After finding integration anchors and running the IntegrateData function, the workflow described above for PCA, clustering, and data visualisation was performed on the integrated dataset to explore ependymal cell heterogeneity. Multiple rounds of differential expression testing were performed on the raw (non-integrated) normalised counts using the functions FindAllMarkers, to identify markers of a cluster compared to all the other cells, and FindMarkers, to identify markers genes that differentiate two specific clusters. A Wilcoxon rank sum test was used for all differential expression testing with a log-fold change threshold of 0.25 and on genes detected in a minimum of 15% of the cells in each cluster. Cell identities were assign based on known markers and informed by marker genes and our own findings for mature lateral cells. Clusters that were not defined by the expression of unique markers were merged with the closest cluster based on pair-wise differential expression analysis and guided by clustering tree visualisations. A phylogenetic tree relating the average cell from each cluster (Figure 2B) was estimated based on a distance matrix constructed in the integrated counts space using Seurat’s BuildClusterTree function.

To identify dissociation-affected cells (Figure S3), we followed a similar approach than van den Brink and colleagues (van den Brink et al., 2017). First, we made a list of dissociated-induced genes in ependymal cells. To do so, we performed a Wilcoxon rank-sum test and to identify genes differentially expressed in the activated lateral cell cluster compared to the lateral cell cluster in homeostasis (setting a log-fold change threshold of 0.25 and using genes detected in minimum 25% of the cells in each cluster). We then used the PercentageFeatureSet function in Seurat to calculate the percentage of reads or UMIs in each cell that mapped to genes in our dissociation-induced genes list. This measure gives an indication of how affected each cell is by the dissociation. We then plotted the distribution of this measure as a histogram for every cell in the dataset and chose a threshold value to define cells with a value that is equal or above this threshold value as dissociation-affected. We chose a threshold cutoff value of 9.3%, or 3 median absolute deviations from the median percentage of the cell’s transcriptome mapping to our list of dissociation-induced genes. Dissociation-affected cells were then marked in a UMAP plot of the integrated dataset.

#### Ependymal cell ageing analysis

Filtered 10x datasets from cells in the spinal cord central canal region of young adult and aged mice were aggregated and processed using Seurat v3.2.2 (Stuart et al., 2019). An initial analysis was performed as described in “scRNA-seq quality control, normalisation, and dimensionality reduction” to take advantage of the larger number of cells in the aggregated dataset in identifying cell clusters and marker genes. We then subset the ependymal cell cluster to study ependymal cell heterogeneity and age-related changes in gene expression and cellular composition. Highly variable features were identified with Seurat’s FindVariableFeatures function using the vst method with default parameters. The top 2,500 highly variable features were used for PCA and significant

PCs were used for graph-based clustering. The same PCs were used as input to UMAP. Clustering was performed at a range of resolutions and clustree was used to visualise relationships between clusters and to identify the most biologically meaningful clustering resolution. After performing differential expression testing using a Wilcoxon rank sum test, cell identities were assigned to each cluster based on known marker genes and data from http://www.mousebrain.org (Zeisel et al., 2018). A cell cluster characterised by the co-expression of markers of astrocytes and ependymal cells was removed after we could not confirm the existence of such cells in the tissue. A 3-cell cluster that expressed choroid plexus genes was also removed from the dataset. After removing potential doublets and contaminating cells, highly variable features, PCA, clustering and visualisation was repeated as described above. Multiple rounds of differential expression testing were performed using the functions FindAllMarkers and FindMarkers. A Wilcoxon rank sum test was used for all differential expression testing with a log-fold change threshold of 0.25 and on genes detected in a minimum of 15% of the cells in each cluster. Cell identities were assign as described in our ependymal cell heterogeneity analysis and clusters that were not defined by the expression of unique markers were similarly merged with the closest cluster based on pair-wise differential expression analysis and guided by clustree. Differential expression analysis was performed to identify age-related changes in gene expression in the ependymal cell population. To do so, age (young or aged) was assigned as cell identities and Seurat’s FindMarkers function with a log-fold change threshold of 0.2 was used to identify genes that were up-or downregulated in cells from aged mice in at least 25% of the cells. To calculate the fraction of each ependymal cluster in each sample, the number of cells from each sample in a given cluster was calculated and normalised to the number of cells per sample. A two-sided Wilcoxon rank sum test was then performed to test for significant differences in the fractions of cluster between young and aged mice using base R. P-values below 0.05 are indicated in Figure 3D.

#### Ependymal cells across the CNS analysis

scRNA-seq data of ependymal cells from across the CNS were obtained from (Zeisel et al., 2018). First, we downloaded a loom file including curated ependymal cell transcriptomes from http://www.mousebrain.org (l6_r4_ependymal_cells.loom). This file was read with Seurat via loomR to extract from which 10x runs these cells were captured and then the raw sequencing reads downloaded for preprocessing and integration with our 10x dataset. All relevant BAM files were converted to FASTQ files using bamtofastq v1.2.0 (10x Genomics) and preprocessed using the same workflow as described above for our 10x data (see “scRNA-seq data preprocessing droplet-based scRA-seq“). Count matrices were loaded to Seurat for further analysis. Data from the following tissues were included: amygdala, hypothalamus, hippocampus, striatum, midbrain, thalamus, pons, medulla, and spinal cord. After removing cells expressing fewer than 500 features and genes detected in fewer than 3 cells, the filtered data were normalised with a scale factor of 10,000 UMIs per cell and then log transformed. Highly variable features were identified with Seurat’s FindVariableFeatures function using the vst method with default parameters. The top 2,200 highly variable features were used for PCA and significant PCs were used for graph-based. The same significant PCs were used as input to UMAP. Clustering was performed at a range of resolutions and clustree was used to visualise relationships between clusters and this, in combination with differential expression analyses, was used to set the most biologically meaningful clustering resolution. Informed by differentially expressed genes, cell clusters were annotated based on known marker genes and data from http://www.mousebrain.org. Clusters assigned to ependymal cells were subset from the dataset and the workflow repeated, from the identification of highly variable genes step to differential expression testing between clusters. Clusters of non-ependymal identity were removed and the workflow repeated again until the dataset contained only ependymal cells. Next, we integrated the Zeisel dataset with our 10x spinal cord ependymal cell dataset from young adult mice using Seurat’s CCA (Stuart et al., 2019). To minimise technical artefacts and challenges due to different Chromium Single Cell 3′ chemistries used to generate the 10x datasets, we kept only genes that were detected in both datasets. The data were merged, normalised with a scale factor of 10,000 UMIs per cell and then log transformed. Highly variable features were identified with Seurat’s FindVariableFeatures function and the vst method. After selecting 2,000 integration features with the function SelectIntegrationFeatures, finding integration anchors and running the IntegrateData function, the standard workflow for PCA, clustering, and data visualisation was performed on the integrated counts. Multiple rounds of differential expression testing were performed on the raw (non-integrated) normalised counts. A Wilcoxon rank sum test was used for all differential expression testing with a log-fold change threshold of 0.25 and on genes detected in a minimum of 25% of the cells in each cluster. Following this, a microglia cluster and a cluster of potential astrocyte and ependymal cell doublets were removed from the dataset and the workflow described above ran again from finding integration anchors and integrating the data, to performing PCA and selecting significant PCs, clustering at a range of resolutions, visualising the data with UMAP, and performing differential expression testing to assign cell identities. Cell identities were assign based on known markers and the tissue of origin as recorded in each cell’s metadata. Clusters not defined by the expression of unique markers were merged with the closest cluster based on pair-wise differential expression analysis and their relationship with other clusters as visualised with clustree.

#### Visualisation

UMAP plots showing gene expression patterns were generated using the R package ggplot2 (Wickham, 2016) and the viridis colour palette (Garnier et al., 2021). Stacked bar plots, bar plots, and boxplots were also generated with ggplot2. All other plots were generated with Seurat.

#### GO enrichment analysis

Marker gene lists were generated using Seurat’s FindAllMarkers or FindMarkers functions and based on the non-parametric Wilcoxon rank sum test with Bonferroni correction. Genes with an adjusted p-value below 0.05 were considered to be differentially expressed. Marker gene lists for each cluster were uploaded as input to g:Profiler for functional profiling (https://biit.cs.ut.ee/gprofiler/gost) (Raudvere et al., 2019). Marker genes lists were ordered from the most significant to the least significant adjusted p-value. GO enrichment analysis was performed using the hypergeometric test and Benjamini-Hochberg FDR correction for multiple testing. GO terms, KEGG, Reactome or WikiPathways with an adjusted p-value below 0.01 were reported.

#### Dual RNAscope and immunofluorescence

##### Mouse samples

Mice aged 21 days (P21), 3 months and 18 months were anaesthetised and then perfused transcardially with ice-cold phosphate buffered saline (PBS). The spinal cord was quickly dissected and fixed in ice-cold 4% paraformaldehyde (PFA) for 2 h at 4 ºC (RT). E18.5 embryos were dissected in ice-cold PBS and their spinal cord fixed in ice-cold 4% PFA for 2 h at 4 ºC. Spinal cord slices from spinal cord slice culture experiments were fixed in ice-cold 4% PFA for 2 h at 4 ºC. After fixing, all samples were washed in PBS and equilibrated in 30% sucrose/PBS overnight at 4ºC before embedding in OCT (Sakura Tissue-Tek) and storing at −80 ºC until processed. Samples were cryosectioned in 10 µm-thick sections and collected in Superfrost Plus slides (Fisher Scientific). RNAscope was performed following the manufacturer’s RNAscope Multiplex Fluorescent Reagent Kit v2 Assay manual (Advanced Cell Diagnostics). Briefly, sections were thawed then covered in Hydrogen Peroxidase for 10 min at RT, washed twice with distilled water, incubated with RNAscope Protease IV for 5 min at RT then washed immediately with PBS. Probes were hybridised for 2 h at 40 ºC in RNAscope’s HybEZ oven then amplified. After washing the slides, we proceeded directly to perform immunofluorescence for GFP. Slides were washed twice 2 min with PBS containing 0.1% Triton X-100 (PBST), blocked for 1 h in blocking buffer (PBST with 10% donkey serum) and incubated with the primary antibody chicken anti-GFP (1:250 in blocking buffer) (Abcam, ab13970) overnight at 4 ºC. Slides were washed 3x 5 min in PBST and then incubated with secondary antibody overnight at 4 ºC. The following day, slides were washed 3x 5min in PBST, counterstained with DAPI and mounted with SlowFade™ Gold Antifade Mountant (Thermo Fisher).

RNAscope probes used were: Mm-*Sntn* (ACD cat.# 574191), Mm-*Vtn* (ACD cat.# 443601), Mm-*Zic1* (ACD cat.# 493121), Mm-*Wnt4* (ACD cat.# 401101), Mm-*Dkk3* (ACD cat.# 400931). Opal™ dyes used were: Opal 520 (cat#FP1487001KT), Opal 570 (FP1488001KT), and Opal 690 (cat#. FP1497001KT) (Akoya Biosciences).

##### Human samples

Fresh-frozen human spinal cord samples (Table S7) from the Netherlands Brain Bank were serially cryosectioned in 10 µm-thick sections and collected in Superfrost Plus slides. RNAscope was carried out following the manufacturer’s RNAscope Multiplex Fluorescent Reagent Kit v2 Assay manual for fresh frozen tissue (Advanced Cell Diagnostics). Briefly, sections were fixed 15 min with fresh, ice-cold 4% PFA and then rinsed 2 times with PBS. Sections were then dehydrated in a series of steps with solutions of increasing ethanol concentration in PBS and then covered with RNAscope Hydrogen Peroxidase for 10 min at RT and washed twice with distilled water. Sections were incubated with RNAscope Protease IV for 2 min at RT then washed immediately with PBS. Probes were hybridised for 2 h at 40 ºC in RNAscope’s HybEZ oven and then amplified. After washing the slides, we proceeded directly to perform immunofluorescence for FOXJ1. Slides were washed twice 2 min with PBST, blocked for 1 h in blocking buffer, and incubated with primary antibody rabbit anti-FOXJ1 (1:50 in blocking buffer) (Sigma-Aldrich, HPA005714) overnight at 4 ºC. Slides were washed 3x 5 min in PBST and then incubated with secondary antibody overnight at 4 ºC. The following day, slides were washed 3x 5min in PBST, counterstained with DAPI and mounted with SlowFade™ Gold Antifade Mountant. RNAscope probes used were: Hs-*SNTN* (ACD cat.# 891411), Hs-*ZIC1* (ACD cat.# 542991), Hs-*ARX* (ACD cat.# 486711), Hs-*PPIB* (ACD cat.# 313901). Opal™ dyes used were: Opal 570 (FP1488001KT), and Opal 690 (cat#. FP1497001KT).

Images from mouse and human samples were acquired with a Leica TCS SP8 confocal laser scanning microscope (Leica Microsystems). All images are shown as maximum intensity projections of acquired z-stacks of about one cell thickness (6-10 µm). Images of the human spinal cord were taken as tiled z-stacks and were stitched together using the stitching algorithm in the Leica Application Suite X (LAS X) software. A minimum of two composite images were taken from each human spinal cord sample and a minimum of 5 images were taken from each mouse sample. Human spinal cord samples, especially those from older individuals, showed relatively high levels of autofluorescence (probably from lipofuscin granules). The probe against the broadly expressed gene *PPIB* was used to (1) control for RNA quality and (2) to differentiate autofluorescence and unspecific probe binding, identified as overlapping *PPIB* and *SNTN* signal, from real signal. Images from outside of the human ependyma were also acquired to get a sense of the level of unspecific binding of the probes. All images were prepared for publication using Fiji (Schindelin et al., 2012).

#### Spinal cord slice culture

Spinal cord slice culture experiments were carried out following Fernandez-Zafra et al. (Fernandez-Zafra et al., 2017) with some modifications. Briefly, 5-month-old mice were anaesthetised and then perfused transcardially with ice-cold, oxygenated aCSF and the spinal cord quickly dissected out in ice-cold aCSF. The thoracic region of the spinal cord was then quickly embedded in 4% low-melting point agarose in PBS (Thermo Fisher) and 350 µm spinal cord slices were cut in ice-cold oxygenated aCSF using a vibratome (Leica VT1200). 3-4 spinal cord slices from each mouse were fixed in 4% PFA for 2 h at 4ºC immediately after vibratome sectioning to serve as the normal/uninjured condition in our experiments. The rest of spinal cord slices were quickly transferred to 30 mm cell culture inserts (Millipore Sigma) previously coated with coated with 10 mg/mL of poly-L-lysine (Millipore Sigma). Each insert with up to 4 spinal cord slices was then was placed in a well of a 6-well plate filled with 1.5 mL of culture medium (50% Neurobasal-A; B-27; 25% heat inactivated horse serum; 25% HBSS; 0.25 mM GlutaMax, 15 mM D-glucose; 15 mM HEPES; and 25 μg/ml penicillin/streptomycin). Every other day, 750 μl of culture medium was replaced with fresh culture medium without disturbing the spinal cord slices or introducing introduce air bubbles. The spinal cord slices were kept in a humidified incubator at 37 °C with a 5% CO2 atmosphere. At the time of collection, cultured spinal cord slices were fixed in ice-cold 4% PFA for 2 h at 4 ºC, then washed in PBS and equilibrated in 30% sucrose/PBS overnight at 4 ºC before embedding in OCT and storing at −80 ºC until processed. Dual RNAscope and immunofluorescence using probes Mm-*Sntn* (ACD cat.# 574191) and Mm-*Mia* (ACD cat.# 498011) and rat anti-Ki67 antibody (Thermo Fisher Scientific, 14-5698-82) was performed as described above for mouse samples.

#### Cell counts and statistical analysis

Images were analysed using Fiji and cells counted with the Cell Counter plugin (Schindelin et al., 2012). For quantification of cells expressing marker genes across ages, ependymal cells were identified based on GFP immunofluorescence and DAPI-stained nuclei in samples from FOXJ1-EGFP spinal cords, and based on location within the central canal walls and nuclear morphology as revealed by DAPI (i.e. small oval nucleus with dense chromatin and multiple nucleoli). All ependymal cells within a 10 µm confocal z-stack were counted. Ependymal cells labelled with one or more RNAscope dots either within the GFP-filled cytoplasm in samples from FOXJ1-EGFP mice or within 2-3 µm of the nucleus were considered positive. For quantification of the fractions of *Sntn* and Ki67-positive cells in cultured spinal cord slices, ependymal cells were identified as *Mia*-expressing cells by RNAscope. All ependymal cells in a single optical slice of a 10 µm z-stack were counted. Ependymal cells with one or more *Sntn* RNAscope within 2-3 µm of the nucleus were considered *Sntn* cells. Ki67-positive cells were identified based on nuclear Ki67 signal.

No statistical method was used beforehand to determine sample size. The investigators were not blinded, and no data points were excluded. Data is represented as mean ± standard deviation. The number of replicates as well as the type of statistical test performed is indicated in each figure legend where relevant.

## Data and code availability

Raw single-cell RNA-sequencing data, count matrices, and metadata are being submitted to Gene Expression Omnibus (GEO) and will be made available upon publication. The code used in this study for scRNA-seq data analysis can be obtained from: https://github.com/aidarodrigo/ependymal_cell

Any additional information required to re-analyse the data reported in this paper is available from the lead contacts upon request.

## Acknowledgements

We thank Rosie Clarke and Arlene Rennie from the School of Life Sciences Flow Cytometry and Cell Sorting Facility for expert FACS assistance, the WBRUTG team for technical assistance, Jeanette Baran-Gale for very helpful and stimulating discussions on scRNA-seq data analysis, Franz Gruber for help with all-things R, Paul Appleton and the Dundee Imaging Facility for microscopy assistance, Edinburgh Genomics for sequencing service in the middle of the COVID-19 pandemic, Geoff Barton for lending A.R.A a desk in Computational Biology. We gratefully acknowledge the Chan Zuckerberg Biohub for support and for sequencing, Foad Green, Steven Chen, Ashley Maynard, Rene Sit, Norma Neff, Spyros Darmanis and members of the Tabula Muris Consortium for technical assistance. This project was supported by the Wellcome Trust (Wellcome Investigator award WT102817AIA to KGS) and a Wings for Life grant to K.G.S. This project and A.R.A. were also supported by the European Union’s Horizon 2020 Marie Skłodowska-Curie grant agreement No. 753812. A.R.A was also supported by the Wellcome ISSF COVID-19 Research Momentum fund (119293). G.A.S. was supported by Wings for Life. C.P.P was supported by the MRC (MC_UU_00007/15). A.P.M. was supported by the ChanZuckerberg Biohub. The confocal microscope used for imaging was purchased with support from Wellcome Trust Multi-User Equipment grant (WT101468).

## Author contributions

A.R.A and K.G.S. conceived and designed the study. A.R.A. generated scRNA-seq data with assistance from G.A.S. A.P.M. generated Smart-seq2 libraries and sequencing data. A.R.A performed data analysis with guidance from C.P.P. A.R.A. and G.A.S. generated RNAscope and immunofluorescence data from mouse and human spinal cord samples. A.R.A. and G.A.S. carried out spinal cord slice culture experiments and RNAscope and immunofluorescence on spinal cord slices. GAS quantified RNAscope and immunofluorescence data. A.R.A. and K.G.S. interpreted the results and A.R.A. with K.G.S. wrote the original draft of the manuscript and all authors read and proofed the final manuscript. K.G.S. and A.R.A. acquired funding.

## Declaration of interests

The authors declare no competing interests.

---

**Figure S1.**
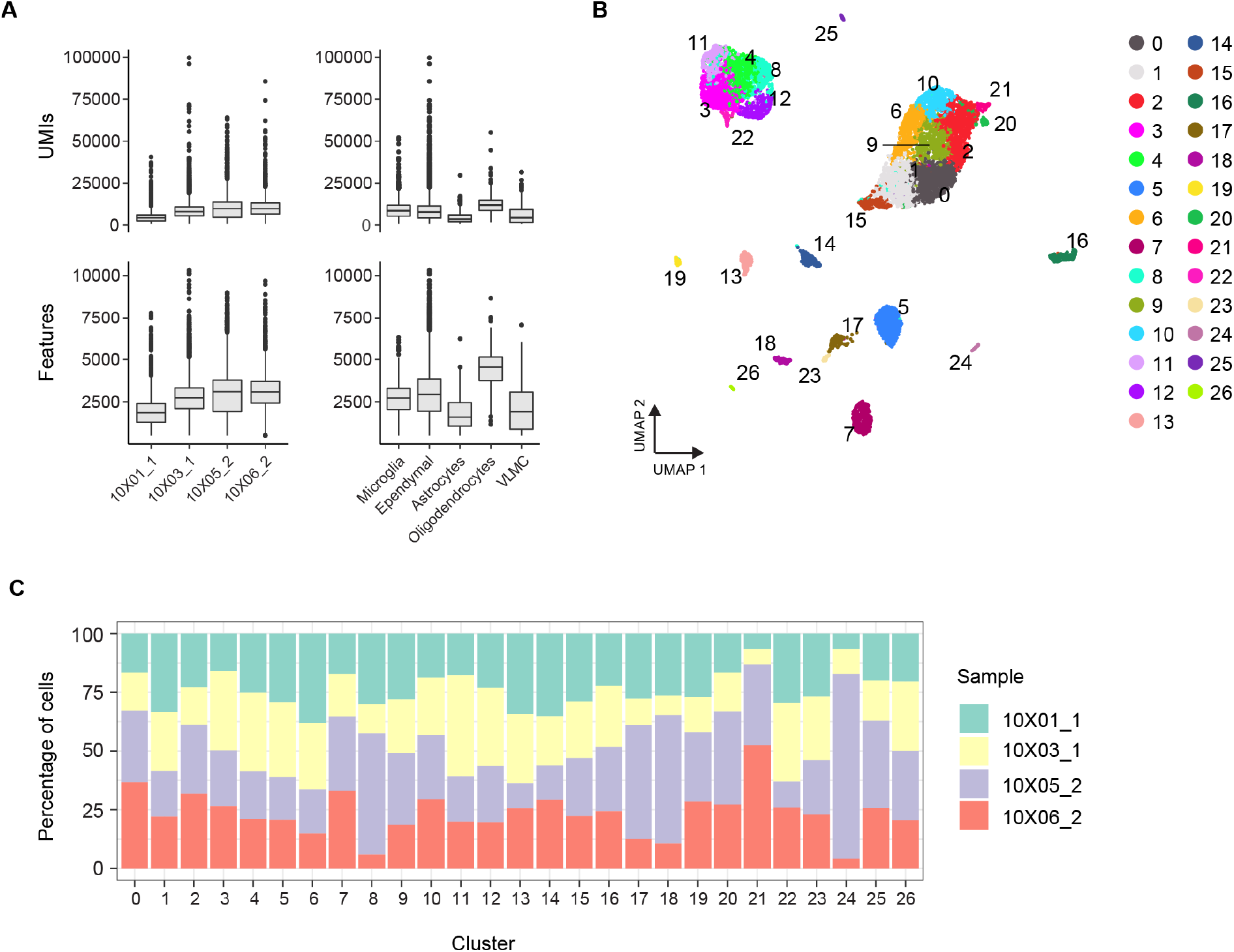
Quality control metrics of the 10x dataset of cells from the central canal region from young adult mice. (A) Box plots showing the distribution of UMI counts and features for all biological samples and for the five most abundant cell types in the dataset. VLMC, vascular leptomeningeal cells. (B) UMAP plot showing further molecular heterogeneity within the clusters in Figure 1C. Cells were initially clustered with Seurat using a clustering resolution of 1.0 and then merged into the 11 cell types represented in Figure 1C. (C) Percentage of cells within each cluster in (B) from each sample, showing contribution of all samples to all clusters.

**Figure S2.**
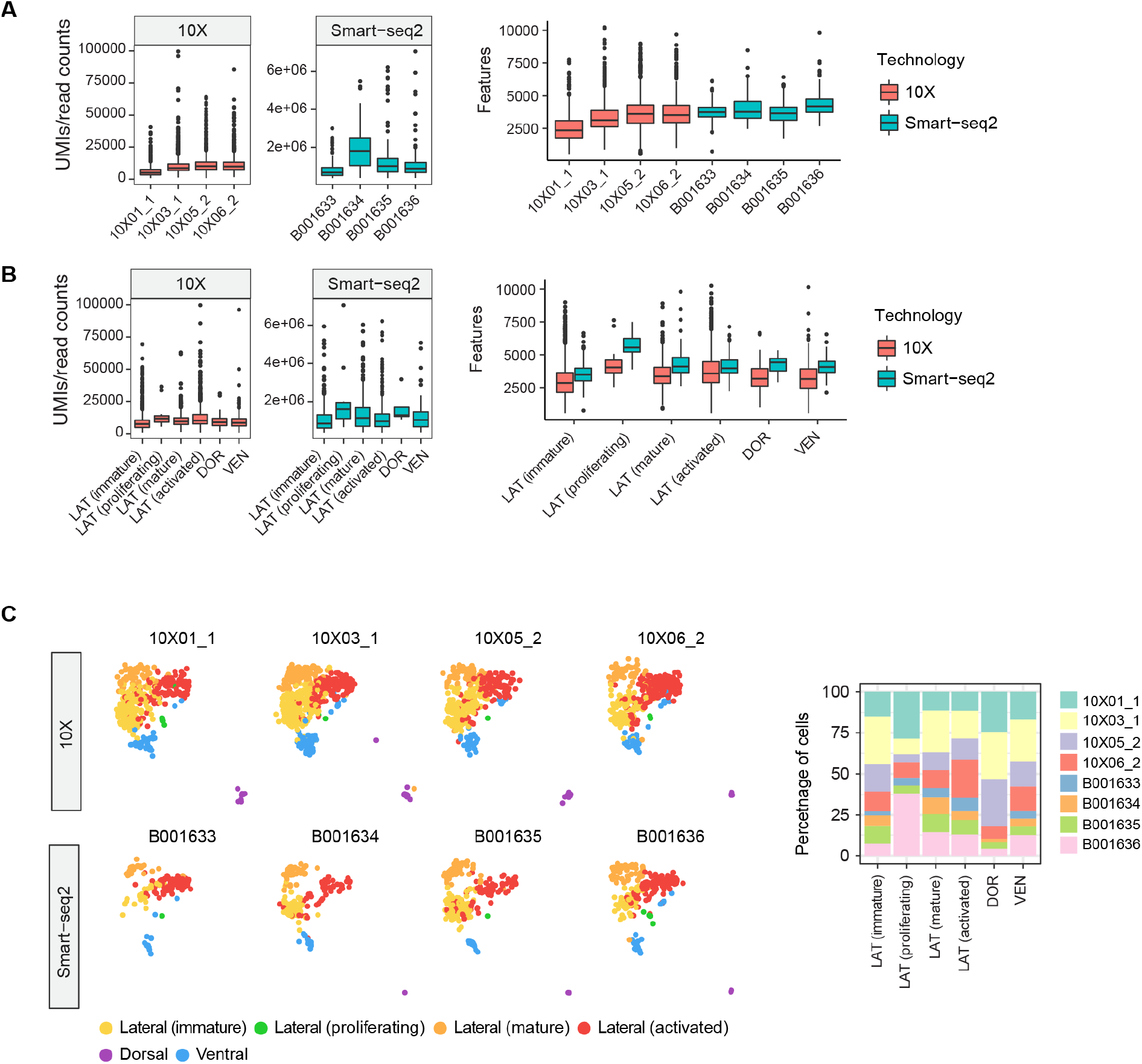
Quality control metrics of the integrated ependymal cell dataset. (A) Box plots showing the distribution of UMIs (10x) or read counts (Smart-seq2) and features for all biological samples and (B) for all the ependymal cell subtypes in the 10x and Smart-seq2 datasets. Note that cells in the Smart-seq2 dataset were sequenced at a much higher depth than cells in the 10x dataset, yet the number of features detected per sample and for each ependymal subtype is relatively similar between technologies. LAT, lateral ependymal cells; DOR, dorsal ependymal cells; VEN, ventral ependymal cells. (C) UMAP plot split by sample showing almost all samples contributed cells to all clusters. Dorsal and proliferating lateral cells lack contribution from one sample. Dorsal and proliferating cells are rare, and this highlights the importance of capturing a large number of ependymal cells in scRNA-seq studies to investigate ependymal cell heterogeneity. Stacked bar plot on the right shows the percentage of cells within each cluster coming from each sample.

**Figure S3.**
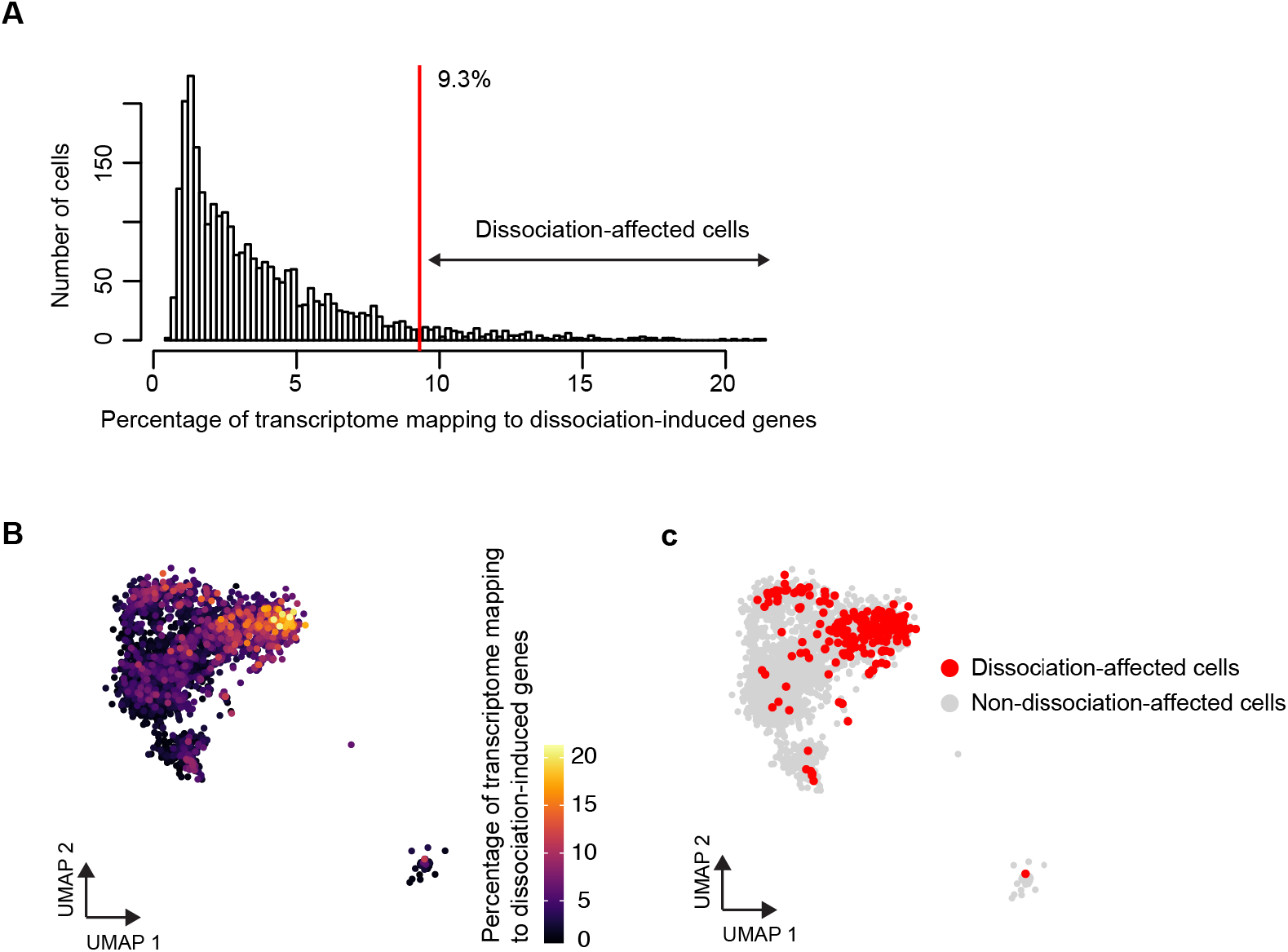
Identification of dissociation-affected ependymal cells. (A) Histogram showing the distribution of the percentage of the transcriptome mapping to dissociation-induced genes in ependymal cells of the integrated dataset. The vertical red line represents a threshold beyond which cells were considered dissociation-affected (median + 3 median absolute deviations). (B) UMAP plot showing the percentage of the transcriptome mapping to dissociation-induced genes for each cell in the dataset. (C) UMAP plot with dissociation-affected cells highlighted. These are cells with more than 9.3% of their transcriptome mapping to dissociation-induced genes as defined in (A)). This threshold to define dissociation-affected cells is conservative because it does not include all cells within the activated lateral cell cluster. However, it does highlight some mature lateral cells as dissociation-affected cells.

**Figure S4.**
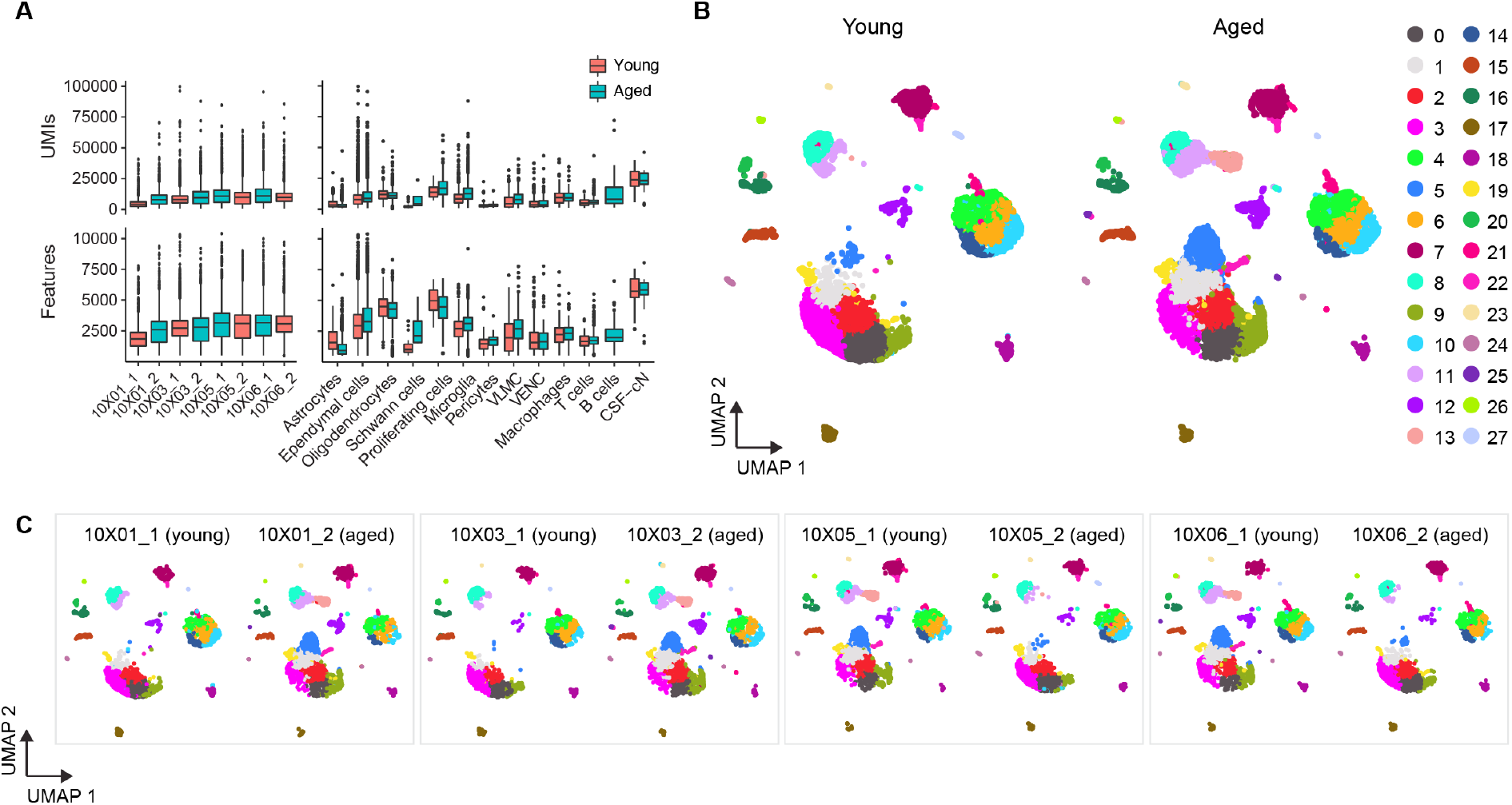
Quality control metrics of the ageing dataset. (A) Box plots showing the distribution of UMI counts and features for all samples (left) and all major cell types (right) identified in the young and aged datasets. VLMC, vascular leptomeningeal cells; VENC, vascular endothelial cells; CSF-cN, cerebrospinal fluid-contacting neurons. (B) UMAP plot showing molecular heterogeneity within the major cell types shown in Figure 3B. For example, a microglia subpopulation expressing *Spp1*, *Lpl*, *Lgals3*, and *Cst7*, was present in low numbers in 3-month-old mice but was clearly more abundant in aged mice (cluster 5). (C) UMAP plots split by sample and colour-coded by cluster as in (B) showing contribution of all samples from the same age to all clusters identified in that age. Samples within the same rectangle were processed on the same day (two mice, one young and one aged were processed in parallel, alternating which sample was dissected and processed first).

**Figure S5.**
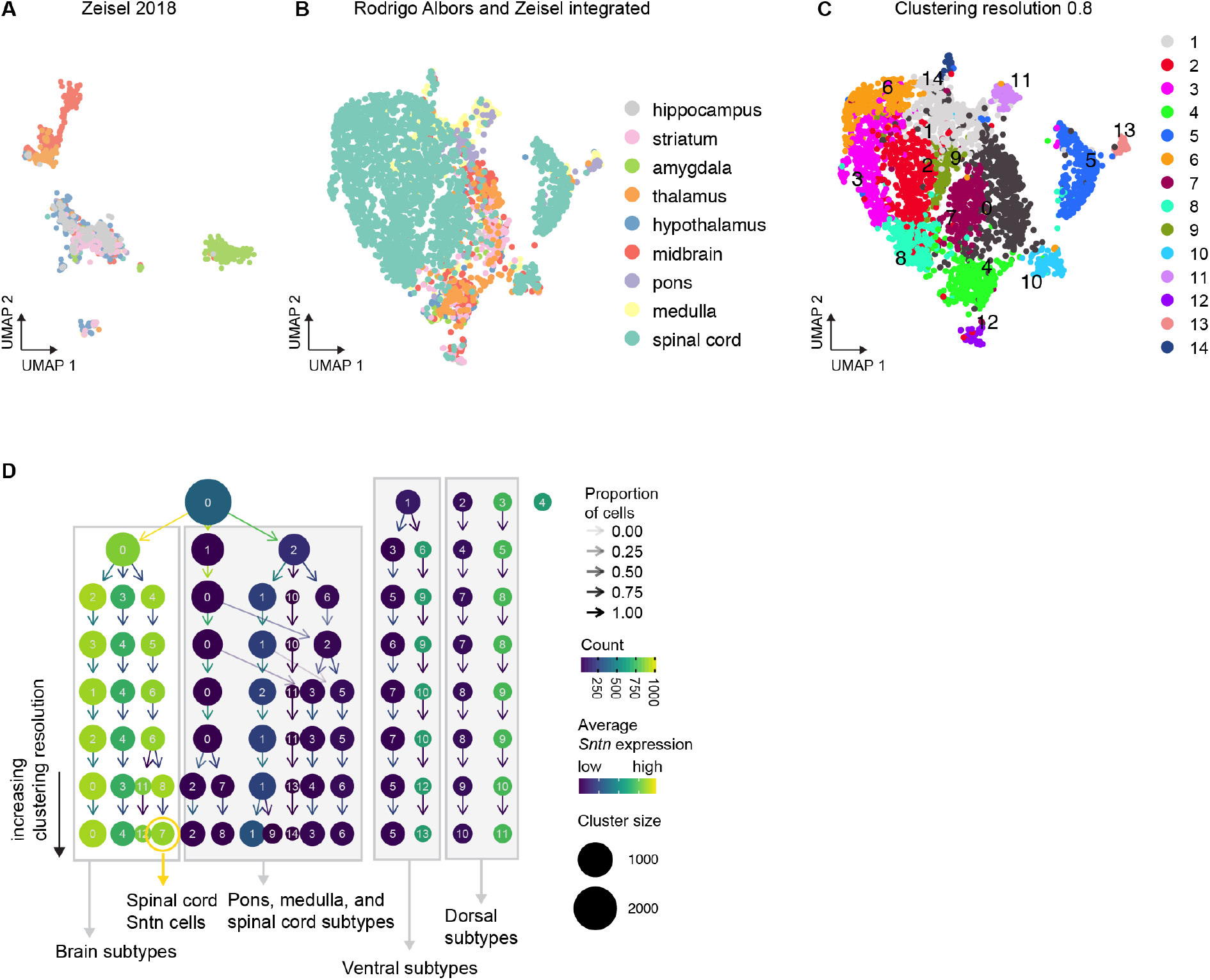
Integration of Zeisel and Rodrigo Albors datasets. (A) UMAP embedding of ependymal cell transcriptomes from Zeisel et al. (Zeisel et al., 2018) coloured by tissue of origin. (B) UMAP embedding of the integrated ependymal cell dataset coloured by tissue of origin. (C) UMAP plot showing molecular heterogeneity within the integrated dataset before merging clusters that were not defined by expression of unique marker genes. Importantly, we were able to identify all spinal cord ependymal cell subtypes and states described earlier, indicating that the integration preserved biological heterogeneity. (D) Clustering tree showing how cells in the integrated dataset split with increasing clustering resolution. The dots represent cluster size and are colour-coded by levels of *Sntn* expression.

**Figure S6.**
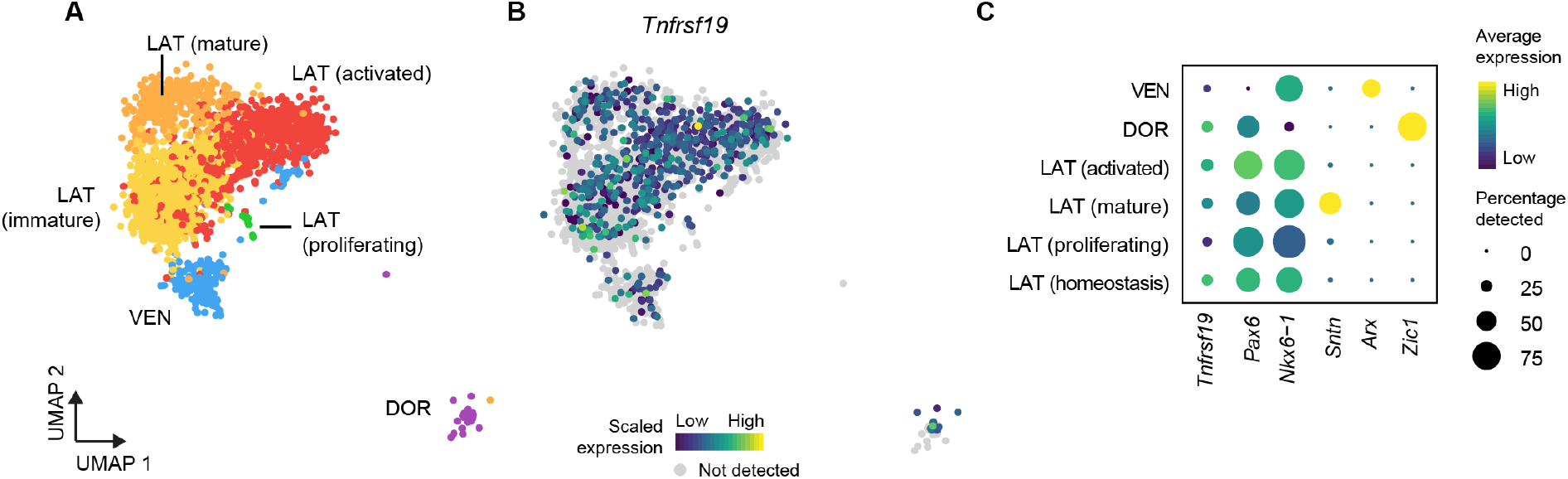
Ependymal cell heterogeneity across the CNS axis. (A) UMAP plot of our integrated spinal cord ependymal cell dataset colour by ependymal cell subtype. (B) UMAP plot showing the expression pattern of *Tnfrsf19*. (C) Dot plot showing the percentage of cells from each ependymal subtype expressing *Tnfrsf19* and a set of marker genes. Dots are colour-coded by average gene expression levels.

## Notes

### Competing Interest Statement

The authors have declared no competing interest.

